# How mutation rate evolves: Gene expression shapes genome-wide mutation rates in *Drosophila melanogaster* and *Caenorhabditis elegans*

**DOI:** 10.64898/2026.11.05.764598

**Authors:** Daniel Phillips, Skirmantas Kriaucionis, Peter Sarkies

## Abstract

Mutation rates vary across the genome, linked to biochemical properties of local DNA sequences. However, mutation rate variation also associates with functional genomic features, raising the possibility that intragenomic mutation rate variation is shaped by adaptive evolution. In particular, recent studies in the model plant *Arabidopsis thaliana* suggested that highly conserved genes under strong purifying selection also have reduced *de novo* mutation rates, suggesting that natural selection may have acted to reduce mutation rate specifically at important genes. However, whether this is true for wider taxa, including animals, remains unknown. Here we repurposed published mutation accumulation datasets from *Drosophila melanogaster* and *Caenorhabditis elegans* to investigate the determinants of *de novo* single nucleotide variant (DNV) rates genome-wide under minimal selection. Loci under stronger purifying selection accumulated fewer mutations, including intronic and synonymous mutations. Various sequence-specific (i.e. GC content) and regulatory features (e.g. chromatin modifications) were strongly correlated with mutation rate, but gene expression best explained the depletion of mutations within strongly conserved loci, even after accounting for correlations amongst variables. Using excision repair (XR)-sequencing data we determined that higher levels of transcription-coupled repair (TCR) were associated with reduced mutation rates, strongly suggesting that TCR lowers mutation rate of highly conserved genes. Furthermore, by comparing orthologs between *C. elegans* and its close relative *Pristionchus pacificus,* we found that altered gene expression across evolutionary time correlated with altered mutation rate. These findings support a model whereby developmental transcription modulates local mutation rates through TCR. A corollary is that adaptive mutation bias may emerge indirectly, downstream of gene regulation evolution, providing a mechanistic framework that reconciles non-random local mutation rate variation with established evolutionary theory.

## 2.0 Introduction

A central principle of population biology is that although evolution can proceed through multiple mechanisms, adaptation is driven by natural selection acting on random mutations. However, germline mutations are far from random in their occurrence. Both mutation rate and mutational spectrum vary systematically within and between genomes, a phenomenon known as *mutation bias*. Almost every property of a genomic site influences its probability of mutating, including local nucleotide context, proximity to telomeres, centromeres and replication forks, repetitive sequence content, propensity to form secondary structures, and dynamic regulatory features such as DNA methylation, chromatin state and transcription (**see Gonzalez-Perez, Sabarinathan, and Lopez-Bigas, 2019 for a review in humans**). But such mutation bias does not contradict the evolutionary concept of “random mutation”, which refers to randomness “*with respect to” fitness (***Futuyma, 2017**), or, equivalently, “the direction of selection” (**Charlesworth, Lande, and Slatkin, 1982**). Following this definition, beneficial mutations should not occur at a higher rate than deleterious ones (**Gregory, 2009**). This concept of mutational randomness arose from classic experiments that refuted the popular view that environmental challenges induce specific adaptive mutations (see **Newcombe, 1949**). Yet, although mutations are spontaneous, the growing evidence for pervasive mutation bias raises a subtler possibility: have genomes evolved to shape mutation biases in ways that ultimately improve fitness?

Genomes are well known to contain mutation “hot” and “cold” spots (**e.g. Nesta, Tafur, and Beck, 2021**). Demonstrating adaptive evolution of local mutation rates, however, requires showing that these regions coincide with genomic features whose mutation is more or less likely to affect fitness. At fine scales, differences in mutability are easily explained by intrinsic sequence properties, which in some cases appears to align with selective pressures. For example, conserved coding regions of essential genes are depleted of mutagenic microsatellite sequences (e.g. **Metzgar, Bytof, and Wills, 2000**; **Lin and Kussell, 2012**). Conversely, eukaryotic gene bodies are flanked by mutagenic short tandem repeat (STR) motifs whose abundance and composition vary with both taxon and gene function, and this has been speculated to increase the evolvability of regulatory regions (**Reinar et al., 2024**). Such preferential localisation of intrinsically stable sequences to regions under purifying selection, and unstable sequences to regions under diversifying or positive selection, is indeed consistent with the idea that selection can shape local mutation bias. More recently, however, even entire gene bodies have been found to harbour fewer *de novo* mutations than surrounding DNA (**Monroe et al., 2022**), while genes themselves are reportedly organised into larger chromosomal domains of relatively high or low mutability that are partly associated with gene function, including in humans (e.g. **Chuang et al., 2004**). Unlike cis-acting intragenic sequence effects, these mutational landscapes suggest trans-acting factors might influence mutation rates across multiple genes simultaneously.

One proposed explanation for such trans-acting effects could be targeting of DNA repair pathways to certain regions of the genome, reducing local mutation rates. Mechanistically, this could involve DNA repair pathways recognising specific transcriptional processes or chromatin modifications. This has been observed in *Escherichia coli*, where loci under purifying selection exhibit reduced *de novo* mutation rates, in part due to transcription-coupled repair (TCR; **Martincorena, Seshasayee, and Luscombe, 2012**). Similarly, in mice, TCR was proposed to reduce germline mutation rates in highly expressed, functionally important genes, although very high expression appeared to increase mutation through transcription-coupled damage (**Xia et al., 2021**). However, these studies used neutral polymorphic variation to assess mutation rate, which cannot be separated from the effects of historical selection. Characterising *de novo* variants (DNVs) therefore requires measuring mutation in the absence of natural selection. Mutation accumulation (MA) experiments overcome this problem by maintaining genetically identical lines at extremely small population sizes (typically one individual for asexual species and a single breeding pair for sexual species). Under these conditions, spontaneous DNVs are largely free from natural selection and accumulate over successive generations, except where mutations cause lethality or sterility. Sequencing these lines after many generations therefore provides an estimate of mutation in the near absence of selection (**Katju and Bergthorsson, 2019**).

Monroe *et al*. (**2022**) analysed data from a large MA study in the plant *Arabidopsis thaliana* and reported reduced mutation rates in highly expressed genes under purifying selection. They argued that this was due to the chromatin modification H3K4me1, enriched at highly transcribed genes, which recruits the DNA repair proteins MSH6 and PDS5, reducing gene-body mutation rates in both *Arabidopsis* and rice (**Quiroz et al., 2024)**. These findings, however, remain controversial. Besides criticisms of how DNVs were identified (**Wang et al., 2023; but see Monroe et al., 2023**), it has been questioned whether the observed depletion of mutations reflects mutation rate itself or residual effects of selection on DNV rates (**Charlesworth and Jensen, 2023**). Additionally, it has been argued on theoretical grounds that the selective advantage of locally reducing mutation rates may be insufficient to overcome the drift barrier in most eukaryotes (**Lynch et al., 2023**), questioning the generalisability of the effect.

In light of these discussions, it is important to test the extent to which the tendency of genes under purifying selection to have intrinsically lower mutation rates is found outside *Arabidopsis*. Here, we utilised MA data for two model animal species, *Drosophila melanogaster* and *Caenorhabditis elegans*, which additionally benefit from rich availability of genomic and epigenomic resources. We show that gene-wise mutation rates are lower at genes under strong purifying selection, with essential and housekeeping loci exhibiting the lowest mutation rates. Although numerous sequence-specific and epigenetic features exhibited strong associations with gene mutation rates, gene expression showed the strongest negative association with mutation rate even after accounting for associations amongst correlated variables. Importantly, using excision-repair sequencing (XR-seq) data to map DNA repair genome wide, we show that reduced mutation rates at highly expressed loci is likely due to increased excision repair targeted to transcribed regions. Crucially, by comparing mutation rate data from *C. elegans* and the closely related nematode *P. pacificus*, we show that changes in developmental expression across evolution correlate with changes in mutation rate, thus suggesting that changes in developmental expression profiles cause mutation rate divergence. Together our results demonstrate that mutation bias protects genes under purifying selection in animals, and provides a mechanistic basis for this effect, suggesting that evolution may have played a role in shaping across animal lineages.

## 4.0 Methods

### 4.1 Variant Data Acquisition and Processing

Characterising *de novo* variants (DNVs) requires measuring mutation in the absence of natural selection. Patterns of genetic variation in natural populations cannot be used directly for this purpose because regions with high or low diversity may reflect differences in selection rather than mutation rate. Mutation accumulation (MA) experiments overcome this problem by maintaining genetically identical lines at extremely small population sizes (typically one individual for asexual species and a single breeding pair for sexual species). Under these conditions, spontaneous DNVs are largely free from natural selection and accumulate over successive generations, except where mutations cause lethality or sterility. Sequencing these lines after many generations therefore provides an estimate of mutation in the near absence of selection. To estimate DNV rates under minimal selection, we used publicly available MA datasets from *C. elegans* (**Saxena et al., 2019; Konrad et al., 2019**), *Pristionchus pacificus* (**Weller et al., 2014; Wang et al., 2026**) and *D. melanogaster* (**Assaf et al., 2017**). *P. pacificus* was included to enable comparisons with *C. elegans*, as both are nematodes. Variants were restricted to single nucleotide polymorphisms (SNPs) on canonical chromosomes and lifted to recent Ensembl assemblies (*D. melanogaster*: BDGP6; *C. elegans*: WS235; *P. pacificus*: El Paco) using Picard LiftOverVcf and the appropriate UCSC chain files. Species-specific snpEff databases were constructed for annotation before variants were checked by chromosome-wide mutation counts and mutation-type frequencies (**Fig. S1-2**) and annotated to canonical transcripts.

### 4.2 Calculating Gene Mutation Rates

Gene-wise mutation rate metrics were calculated from the species-specific DNV datasets. Analyses were restricted to gene-body mutations, excluding variants annotated by snpEff as `upstream_gene_variant`, `downstream_gene_variant` or `intergenic_region`. Variants were merged with species-specific GTF annotations using canonical transcript IDs to obtain gene-level features, including gene-body length and nucleotide composition (AT/GC). For each protein-coding gene, the overall mutation rate was calculated as (mutation count + 0.5) divided by gene-body length. Substitution-specific mutation rates were calculated for each mutation class (T>A, C>G, C>T, C>A, T>G and T>C) as (class-specific mutation count + 0.5) divided by the number of corresponding reference nucleotides within the gene body. Synonymous variants were identified using snpEff annotations. The number of synonymous sites (S) for each canonical coding sequence was estimated using the Li–Comeron method implemented in the `seqinr::kaks` function. Only complete coding sequences (length divisible by three and lacking internal stop codons) were retained. Synonymous mutation rates were then calculated as (synonymous mutation count + 0.5) divided by S. Intronic variants were identified from their snpEff annotations, and intronic mutation rates were calculated as (intronic mutation count + 0.5) divided by total intronic length. All analyses were restricted to protein-coding genes.

### 4.3 Estimating Evolutionary Constraint and Association with Mutation Rate

Selective constraint was estimated as the ratio of nonsynonymous (dN) to synonymous (dS) substitution rates between each focal species and a closely related sister species (*C. elegans*–*C. remanei*, *D. melanogaster*–*D. simulans*, *P. pacificus*–*P. exspectatus*). Orthologues were identified using a reciprocal best-hit (RBH) approach. Protein sequences were aligned with the Needleman–Wunsch algorithm, converted to codon alignments using PAL2NAL, and dN and dS estimated using the Comeron method (**Comeron, 1995**) implemented in the dNdS function. Corrupt or incomplete coding sequences were excluded. Associations between log(dN/dS + 0.5) and log(mutation rate) were tested using both Pearson’s and Spearman’s correlations. Genes were also stratified into equal-sized dN/dS quantiles using ggplot2::cut_number. To minimise random assignment of tied dN/dS values, a small amount of random noise (±1 × 10⁻⁶) was added before binning. Directional associations between dN/dS quantile and mutation rate were assessed using two-sided Jonckheere.

### 4.4 Bulk RNA-Sequencing Data Acquisition and Association with Mutation Rate

Bulk RNA sequencing (RNA-seq) datasets from multiple tissues, developmental stages, and germline cell types were downloaded from public databases and harmonised with mutation rate data to associate gene expression and mutation rate. For *D. melanogaster,* somatic expression datasets were downloaded from modENCODE, subsetting to “modENCODE_mRNA-Seq_tissues” for (n=50) tissue transcriptomes, and “modENCODE_mRNA-Seq_development” for (n=12) embryonic timepoints (**Graveley et al., 2011**). Expression data from the apical, middle, and basal testis regions was acquired from Vedelek et al. (**2018**), corresponding to germinal stem cells and mitotic spermatogonia, primary meiotic spermatocytes and spermatids, and elongated spermatids and mature spermatozoa, respectively. For *C. elegans*, expression data for four somatic tissues (muscle, neurons, hypodermis, intestines) and total (primarily hermaphrodite) germline were obtained from the Ahringer lab *C. elegans* regulatory atlas v0.5.4 (**Serizay et al., 2020**). *C. elegans* embryonic gene expression data was downloaded from NCBI database spanning 50 developmental timepoints (**Hashimshony et al., 2015**). Additional expression datasets from oogenic and spermatogenic gonads were downloaded from (**Ortiz et al., 2014**) and library-size normalization was performed using DESeq2 size-factor estimation on the full count matrix. Within-condition replicates were averaged to obtain a per-condition expression value. *P. pacificus* embryonic gene expression was downloaded from the gitHub repository associated with (**Reich et al., 2025**) spanning six developmental time points and gene names were converted to WBGene IDs using the El paco GTF.

Expression data units varied across species (Dmel = [Tissue and Embryogenesis is RPKM, Testis is FPKM], Cel = [Tissue is TPM, Embryogenesis is CEL-Seq normalized abundance, Oogenic and spermatogenic gonads are DESeq2 normalised counts], Ppa = DESeq2-normalised counts), but were processed similarly as log2(expression + 1). Duplicate gene identifiers within each sample were averaged. For each species and developmental timepoint, Pearson’s and Spearman’s correlations were calculated between log₂(expression + 1) and log(mutation rate). Within each transcriptome, genes were binned by their expression levels into six different groups; 0 if they have zero expression, and 1-5 approximately equal sized quantile bins of increasing expression using ntile(). An association between expression bin and mutation rate was tested using two-sided Jonckheere. Statistical tests were also repeated only using genes with at least one mutation. For some cross-tissue visualisations, correlation coefficients were averaged within tissues (e.g. embryo) to improve clarity.

### 4.5 Global Run-On Sequencing Data Analysis

Strand-specific Global Run-On sequencing (GRO-seq) data were obtained for wild-type *C. elegans* embryos (**Kruesi et al., 2013**) and untreated *D. melanogaster* S2 cells (**Fuda et al., 2015**). For *C. elegans*, replicate-averaged, RPKM-normalised WIG tracks were converted to bedGraph and lifted from WS230 to WBcel235 (ce11). For D. melanogaster, normalised bedGraph signals were converted to absolute values to account for negative-strand encoding, lifted from dm3 to dm6, and averaged across two replicates separately for each strand, treating missing intervals as zero. Sense and antisense transcription were assigned according to annotated gene orientation. Mean GRO-seq signal was calculated across gene bodies (TSS to TTS) using base-pair-weighted coverage.

### 4.6 Clustering of Developmental Time Series

Unscaled gene expression profiles were clustered separately for each species to identify groups of genes with similar developmental expression dynamics and assess their relationship with mutation rate. Genes with missing expression values at any developmental timepoint were excluded. K-means clustering was performed using the `ComplexHeatmap` package with visually selected cluster numbers (k = 7 for *C. elegans*, k = 6 for *P. pacificus* and k = 9 for *D. melanogaster*), using 100 random starts to improve cluster stability. Mean expression trajectories were calculated for each cluster and plotted with 95% confidence intervals. Differences in gene mutation rates among clusters were assessed using Kruskal–Wallis tests.

### 4.7 Epigenetic and XR-Seq Data Acquisition and Association with Mutation Rate

Epigenetic covariates were compiled from multiple sources (**Table S1**) in bigWig or BedGraph format. bigWig files were converted to BedGraph using `bigWigToBedGraph`, while replicate bigWig files were merged using `bigwigCompare` (mean operation, 25 bp bins). Datasets mapped to superseded genome assemblies were lifted over to current reference builds (*D. melanogaster*: BDGP6; *C. elegans*: WS235) using UCSC LiftOverVcf and the appropriate chain files, as described above. A key dataset comprised excision repair sequencing (XR-seq) data for *D. melanogaster* (**Deger *et al*., 2022**) and *C. elegans* (**GSM8595214; Kose *et al*., 2025**), which measures genome-wide nucleotide excision repair at single-nucleotide resolution. XR-seq performed in XPC knockout mutants following UV exposure specifically quantifies transcription-coupled repair (TCR) of cyclobutane pyrimidine dimers (CPDs), as global genome repair is inactive.

Epigenetic data were imported into R, converted to genomic ranges, and intersected with gene-body coordinates. For each gene, the mean signal of each epigenetic mark was calculated across all overlapping regions. Associations between mean gene-body signal and mutation rate were assessed using Pearson’s and Spearman’s correlations. Genes were also grouped into five approximately equal-sized quantiles based on mean epigenetic signal, and directional trends in mutation rate across quantiles were tested using two-sided Jonckheere tests. To avoid arbitrary assignment of tied values, a small amount of random noise (±1 × 10⁻⁶) was added before binning. To examine spatial distributions of epigenetic marks, signal profiles were summarised across scaled gene bodies with 1 kb flanking regions using the `normalizeToMatrix` function from the `EnrichedHeatmap` package. Mean enrichment profiles and standard errors were calculated separately for genes grouped by mutation-rate or expression quantiles.

To evaluate the combined effects of multiple predictors, least absolute shrinkage and selection operator (LASSO) regression was performed using the `glmnet` package. For each species, genes with complete data were randomly split into training (80%) and test (20%) datasets, and predictor variables were standardised before model fitting. Model performance was evaluated on the test set using the coefficient of determination (R²) and root mean squared error (RMSE). Predictor importance was assessed from the magnitude and direction of LASSO coefficients and compared with univariate Spearman correlations. Additional gene-level covariates included log₂(expression + 1) and GC content.

### 4.8 Comparative Transcriptomic Analysis of *C. elegans* and *P. pacificus*

To investigate whether evolutionary changes in gene expression or epigenetic features are associated with mutation-rate divergence, one-to-one orthologues between *C. elegans* and *P. pacificus* were identified using `OrthologR::dNdS`, as described above. Gene-level features, including mutation rate, log₂(expression + 1), GC content, mean epigenetic mark signal and expression quantile, were annotated to each orthologue. To facilitate comparisons between species with different distributions, each feature was converted to a within-species percentile rank among orthologues with complete data for that feature. Divergence was then calculated as Δ (*P. pacificus* − *C. elegans*) for each feature and for mutation rate. Associations between Δ(feature) and Δ(mutation rate) were assessed using Spearman’s rank correlations. For epigenetic features, orthologues were additionally grouped into five approximately equal-sized quantiles based on Δ(feature), and directional trends in Δ(mutation rate) were tested using two-sided Jonckheere tests. To avoid arbitrary assignment of tied values, a small amount of random noise (±1 × 10⁻⁶) was added before binning. Orthologues showing the largest increases or decreases in relative percentile (Δ(feature) > 0.6) were also compared with an equal number of randomly selected orthologues exhibiting minimal change (|Δ(feature)| < 0.2) using Wilcoxon rank-sum tests. For gene expression, additional analyses used discrete within-species expression quantiles. Expression bins (0–5) were recalculated separately within each species among the retained one-to-one orthologues, with zero-expression genes assigned to bin 0 and expressed genes divided into five approximately equal-sized quantiles. Changes in expression bin between species were then determined for each orthologue pair. Mutation-rate divergence was compared between orthologues that increased or decreased expression bin and those remaining in the same bin using two-sided Wilcoxon rank-sum tests. The effect of the magnitude of expression change was also evaluated by comparing different quantile-step changes. All tests were repeated using a subset of orthologs with complete expression and dN/dS data with complementary tests using dN/dS as the dependent variable, to determine if divergence in mutation rates linked to expression divergence depends on shifts in the dN/dS.

### 4.9 Housekeeping Genes and Essentiality

Housekeeping and essential gene annotations were obtained from published sources. For *C. elegans*, housekeeping genes were defined as the low-Gini gene set and essential metabolic genes as the high- and medium-confidence RNAi phenotype classes (G1 and G2) reported by Joshi et al. (**2022**). For *D. melanogaster*, essential genes were inferred from FlyBase allele phenotype annotations (**Larkin et al., 2020**) by identifying genes with alleles annotated as lethal and mapping these alleles to their corresponding gene.

### 4.10 Data Visualisation

All data visualisation was produced using ggplot in R. Boxplot statistics follow ggplot2:: geom_boxplot() default settings. Namely, the median value as the central line, the first and third quartiles as the lower and upper edges of the box, and the whiskers extend to the most extreme values within 1.5x the interquartile range (IQR).

## 5. Results

### 5.1 Mutation Rates Reflect Evolutionary Divergence and Conservation

To characterise patterns of DNV across eukaryotes, we analysed published MA datasets from *D. melanogaster* and *C. elegans*. Gene-body mutation rates showed substantial genome-wide variation (**Fig. S3**).

**Table 1.** Total number of unique *de novo* single nucleotide variants (DNVs) identified in previously published mutation accumulation experiments in *D. melanogaster* and *C. elegans*. Variants are classified according to whether they occur within the gene body of at least one protein-coding gene or in intergenic regions.

| **Species** | **Total variants** | **Coding gene body** | **Intergenic** |
|---|---|---|---|
| *D. melanogaster*<br>Assaf et al., 2017 | 3,181 | 2,047 | 1,134 |
| *C. elegans*<br>Saxena et al., 2019<br>Kondrad et al., 2019 | 5,033 | 3,051 | 1,982 |

Previous studies in *Escherichia coli* (**Martincorena, Seshasayee, and Luscombe, 2012)** and *Arabidopsis thaliana* (**Monroe et al., 2022**) reported reduced DNV rates in genes under stronger purifying selection. To test whether this pattern extends to animals, we estimated dN/dS for each gene using orthologues from closely related sister species, with dN/dS values <<1 indicating stronger purifying selection. Gene-body mutation rates were positively correlated with dN/dS in both species (***D. melanogaster*: Spearman’s ρ = 0.236, P < 2.2 x 10^-16^; *C. elegans*: ρ = 0.14, P < 2.2 x 10^-16^**), and these associations remained consistent across substitution classes and when analyses were restricted to genes that accumulated at least one mutation (**Fig. 1a**). Crucially, associations also remained significant for synonymous (***D. melanogaster*: Spearman’s ρ = 0.128, P < 2.2 x 10^-16^; *C. elegans*: ρ = 0.12, P < 2.2 x 10^-16^**) and intronic (***D. melanogaster*: Spearman’s ρ = 0.23, P < 2.2 x 10^-16^; *C. elegans*: ρ = 0.11, P = 6.116 x 10^-13^**) mutation rates (**Fig. 1a**), which are less likely to experience residual selection. When genes were grouped into dN/dS quantiles, mutation rate increased with dN/dS (***D. melanogaster*: Jonckheere P < 0.001; *C. elegans*: P < 0.001**), although the lowest dN/dS bin showed a slight elevation relative to a linear trend (**Fig. 1b-c**). Essential genes had lower overall, synonymous and intronic mutation rates than non-essential genes in both species (**Fig. 1d**), with the exception of *C. elegans* intronic mutation rates, which showed the opposite trend, although the paucity of introns in *C. elegans* meant that few variants were in this category (**Fig. 1d**). Housekeeping genes in *C. elegans* also showed reduced overall, synonymous and intronic mutation rates (**Fig. 1e**).

**Figure 1.**
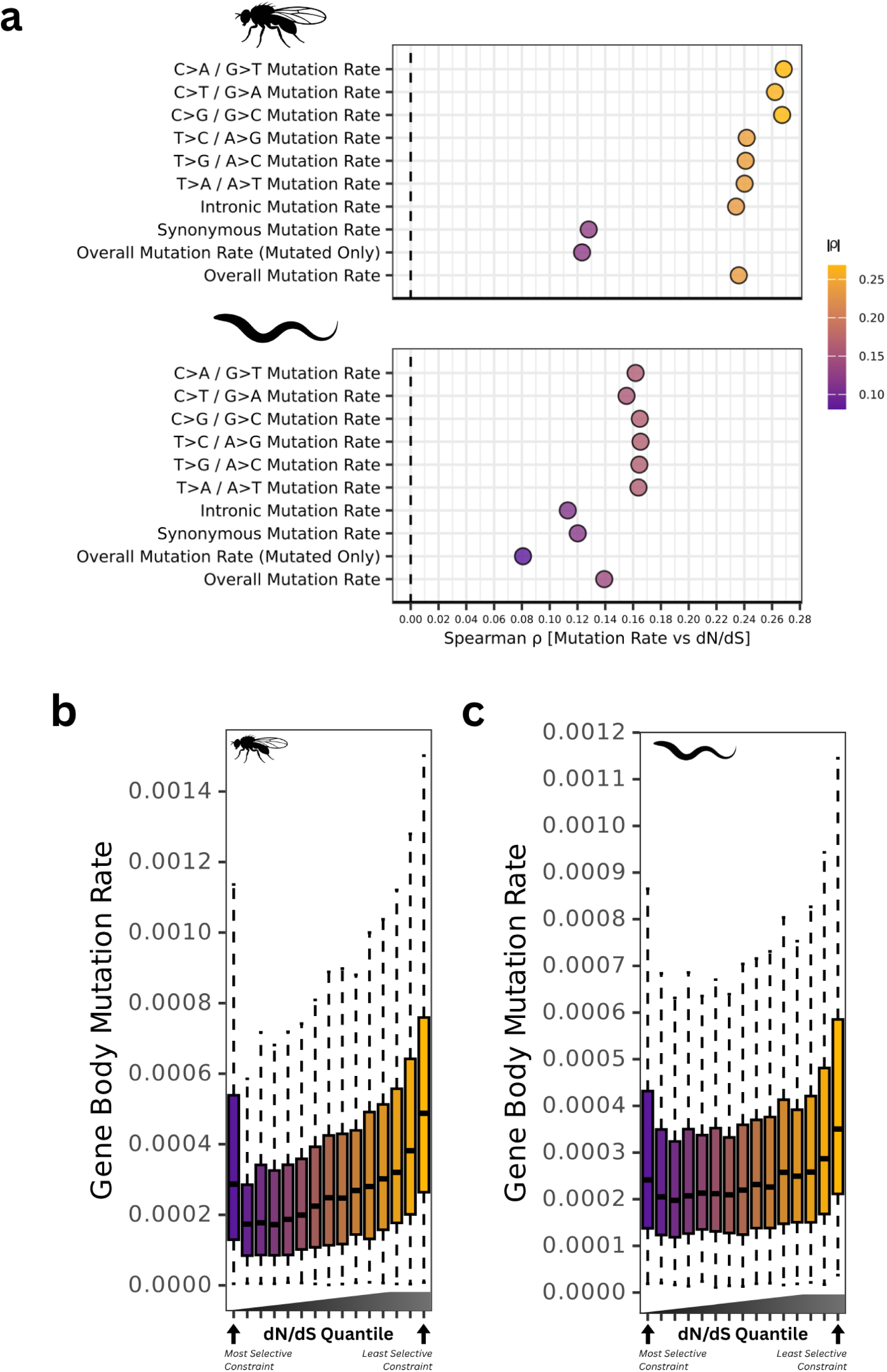
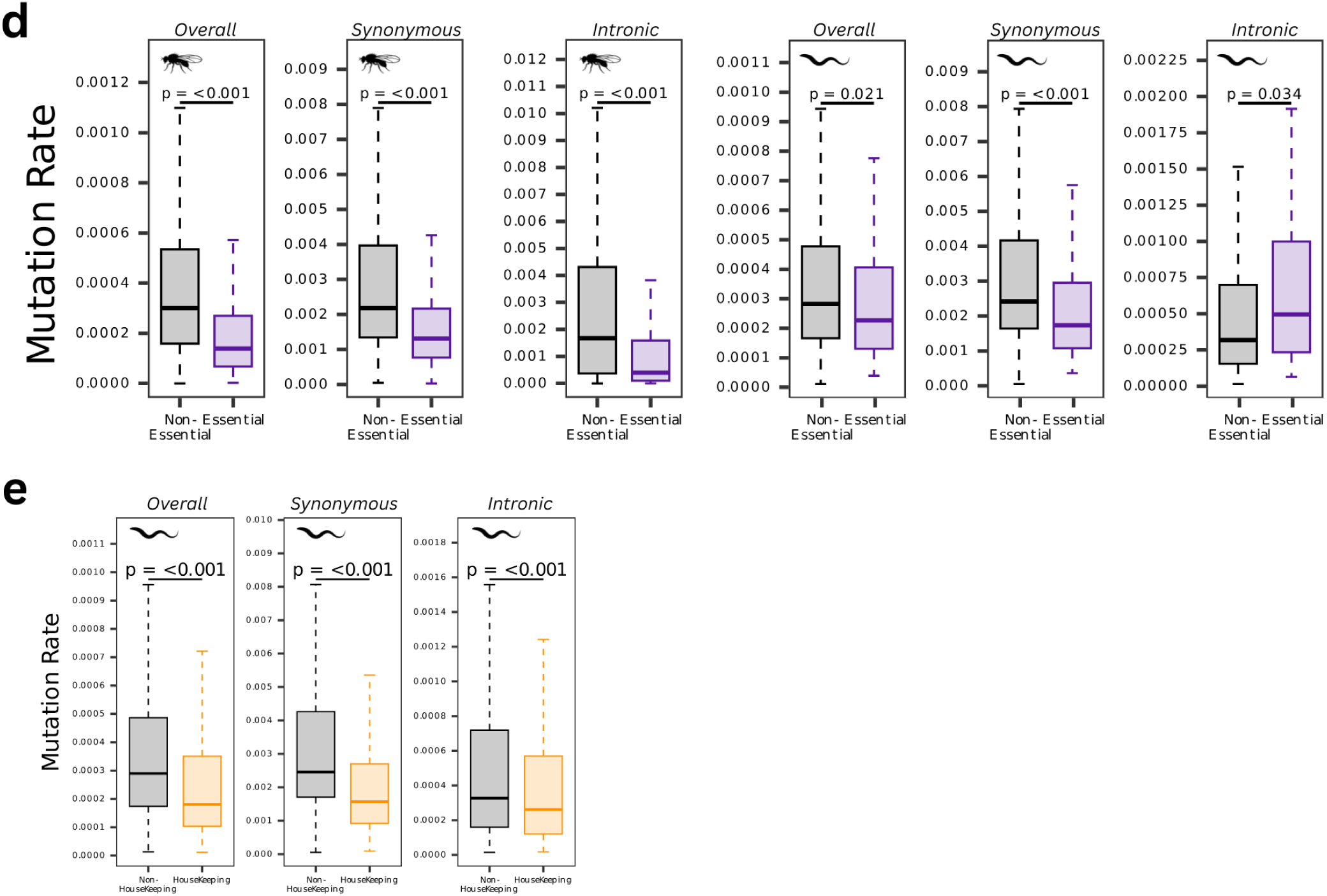
Mutation rates reflect patterns of evolutionary constraint. **(a)** Spearman correlation coefficients between log(mutation rate) and log(dN/dS + 0.5) for each mutation class and species. Point size represents –log₁₀(adjusted P value), and colour indicates correlation coefficient. **(b,c)** Gene-body mutation rates across dN/dS quantile bins in *D. melanogaster* and *C. elegans*. **(d)** Mutation rates in essential and non-essential genes. **(e)** Mutation rates in housekeeping and non-housekeeping genes in *C. elegans*. Reported P values result from Wilcoxon tests.

### 5.2 Developmental Gene Expression is Associated with Mutation Rate

One possible explanation for why essential and housekeeping genes might have lower mutation rate could be their higher expression. To test this, we next asked whether gene regulation was associated with variation in mutation rate, specifically gene expression. To assess the relationship between gene expression and mutation rates in an unbiased manner, we downloaded bulk RNA-seq datasets from various somatic, embryonic, and germline sources. Across all datasets and both species, gene expression was negatively associated with mutation rate, with the exception of the testis in *D. melanogaster* (**Fig. S4a**), which instead exhibited a positive association (**Fig. 2a-b**). There were nonetheless large differences in the strength of the correlations and their consistency across species. Namely, in *D. melanogaster* the strongest mean negative associations were detected in the central nervous system (**ρ = -0.39, P < 2.2 x 10^-16^; Fig. S4b)** and head (**ρ = -0.36, P < 2.2 x 10^-16^; Fig. S4c)**, followed by embryo (**ρ = -0.27, P < 2.2 x 10^-16^**; **Fig. 2a**). In *C. elegans*, oogenic (**ρ = -0.28, P < 2.2 x 10^-16^; Fig. S4d)** and spermatogenic gonads (**ρ = -0.22, P = P < 2.2 x 10^-16^; Fig. S4e**) exhibited the strongest correlation, also followed by embryo (**ρ = -0.18, P < 2.2 x 10^-16^; Fig. 2b**). Because RNA-seq measures steady-state transcript abundance, which is influenced by both transcription and RNA stability, we investigated whether the observed associations with mutation rate persisted when transcriptional activity was measured more directly. To this end, we used Global Run-On sequencing (GRO-seq; **Core, Waterfall, and Lis, 2008; Kruesi et al., 2013; Fuda et al., 2015**), which measures nascent transcription by transcriptionally engaged RNA polymerases in a strand-specific manner. Mean sense-strand GRO-seq signal across gene bodies was significantly negatively correlated with mutation rate in both *D. melanogaster* (Spearman’s ρ = −0.22, P < 2.2 × 10⁻^16^) and *C. elegans* (ρ = −0.24, P < 2.2 × 10⁻^16^). This association was maintained when loci were binned into equal sized quantile groups based on their expression level, indicating the trend is consistent across the entire transcription level distribution (**Fig S5a-d**). These findings demonstrate that the inverse association between transcription and mutation rate is not restricted to steady-state RNA abundance, but is also observed when measuring nascent transcriptional activity. We next chose to focus on the role of gene expression across embryogenesis, due to embryogenic expression being strongly associated with mutation rate in both species whilst involving major transcriptional reprogramming events and the establishment of cell lineages.

**Figure 2.**
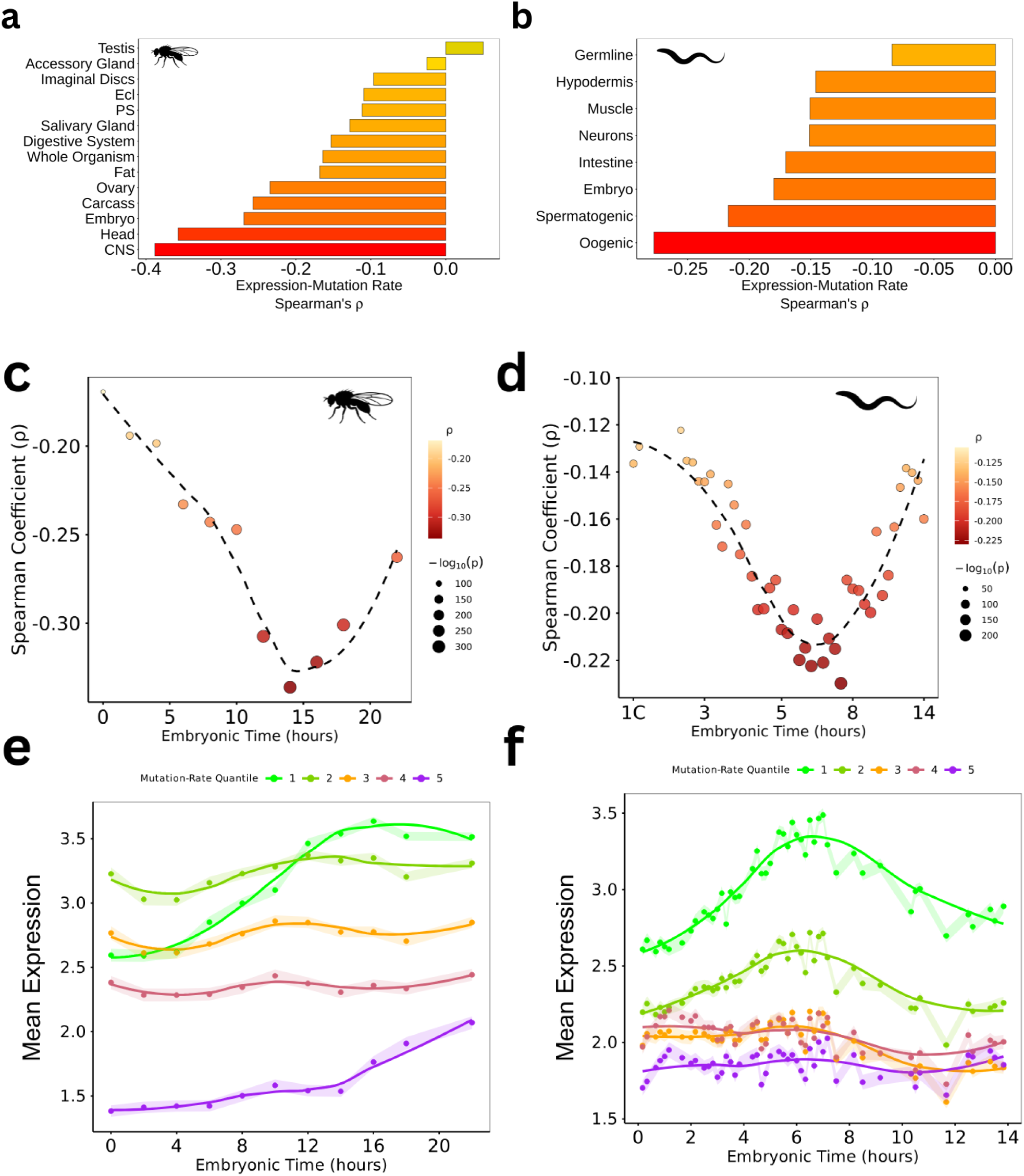
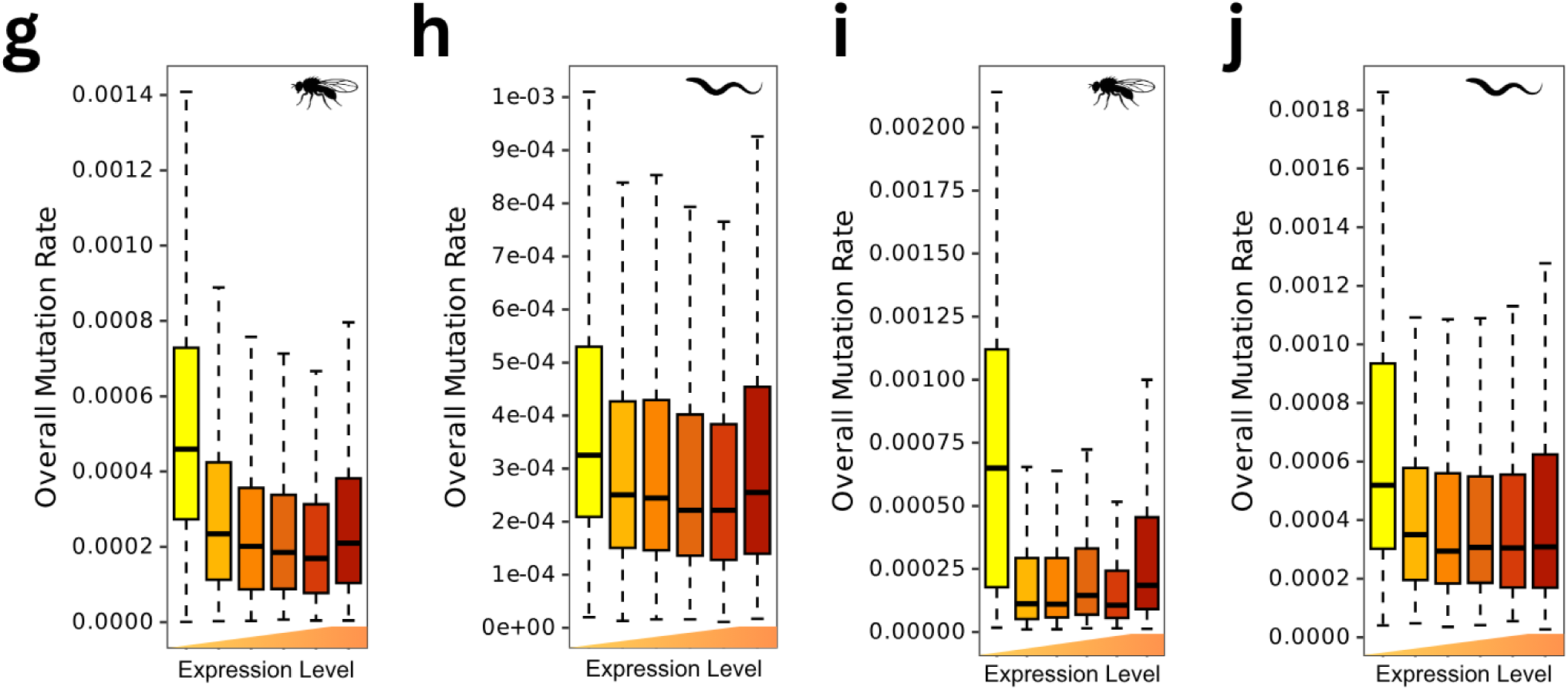
Mutation rates are negatively associated with developmental gene expression. **(a,b)** Bar plots showing the Spearman correlation coefficients between gene expression and mutation rate in different somatic, embryonic, and germline tissues in **(a)** *D. melanogaster* and ***(b)*** *C. elegans*. **(c,d)** Spearman correlation coefficients between protein-coding gene expression and mutation rate across developmental time in *D. melanogaster* **(c)** and *C. elegans* **(d)**. Colours indicate Spearman correlation coefficient while point size represents −log^10^(adjusted P value). **(e,f)** Mean developmental expression profiles of protein-coding genes grouped into five mutation-rate quintiles (1 = lowest mutation rate, 5 = highest mutation rate) in *D. melanogaster* **(e)** and *C. elegans* **(f)**. Shaded ribbons indicate 95% confidence intervals. **(g,h)** Mutation rates of genes grouped into expression quintiles according to late (14-16hr) developmental expression in *D. melanogaster* **(g)** or mid (430min) developmental expression in *C. elegans* **(h)**. **(i,j)** As in **(g,h)**, but restricted to genes accumulating at least one mutation during mutation-accumulation experiments. Box plots show the median and interquartile range, with whiskers extending to 1.5× the interquartile range.

Embryonic expression was negatively correlated with mutation rate at every timepoint in both species, with the strongest associations in mid-late embryogenesis (**Fig. 2c-d**). The same pattern was observed for synonymous, intronic and substitution-specific mutation rates (**Fig. S6a-b**). Genes in the highest mutation-rate quantile had the lowest expression, whereas genes in the lowest mutation-rate quantile peaked in expression mid-late development (**Fig. 2e-f**). At the developmental timepoints with the strongest associations, dividing genes into six expression quantiles showed that step-wise increases in expression were associated with step-wise reductions in mutation rate (***D. melanogaster*: Jonckheere P < 0.001; *C. elegans*: P < 0.001**; **Fig. 2g-h**), a pattern which persisted when only considering genes with at least one mutation (***D. melanogaster*: Jonckheere P = 0.002; *C. elegans*: P < 0.001**; **Fig. 2i-j**). However, dividing genes into 16 expression quantiles revealed a non-linear pattern: the highest mutation rates occurred among both the most lowly expressed and most highly expressed genes (**Fig S7a-b**).

To investigate this relationship using an independent approach, we performed k-means clustering to group genes according to their expression profiles across development (**Fig. 3**). Mutation rates differed significantly among expression clusters in both species (***D. melanogaster*: Kruskal–Wallis, P < 2.2 x 10^-16^; *C. elegans*: P < 2.2 x 10^-16^; Fig. 3a–d**). Genes with little or no expression throughout development and under least constraint (cluster #1 in both species) and those with the highest expression and most constraint (cluster #9 in *D. melanogaster* and #7 in *C. elegans*) had the highest mutation rates, whereas clusters with peak expression during mid-embryogenesis and high constraint (cluster #4 in both species) had the lowest (**Fig. 3a-d**). The high expression-mutation rate clusters were overwhelmingly enriched for terms related to translation and ribosome in both species, whereas the low-expression high-mutation rate clusters were enriched for terms linked to signal transduction and environmental sensing (**Fig S8a-d**). The low mutation rate (#4) clusters were enriched for developmental functions, including morphogenesis, specifically neuronal development, as well as DNA binding and RNA polymerase II regulation in *D. melanogaster*, and primarily transmembrane transport as well as transcriptional and metabolic regulation in *C. elegans* (**Fig. S7e-f**). Gene set enrichment analysis (GSEA) directly on mutation rate distributions produced similar results (**Fig. S9**). GSEA directly on mutation rate distributions additionally revealed an enrichment of genes related to chromatin components and regulation as mutation rate increases in both species, and genes related to sexual reproduction in *D. melanogaster* (**Fig. S9**).

**Figure 3.**
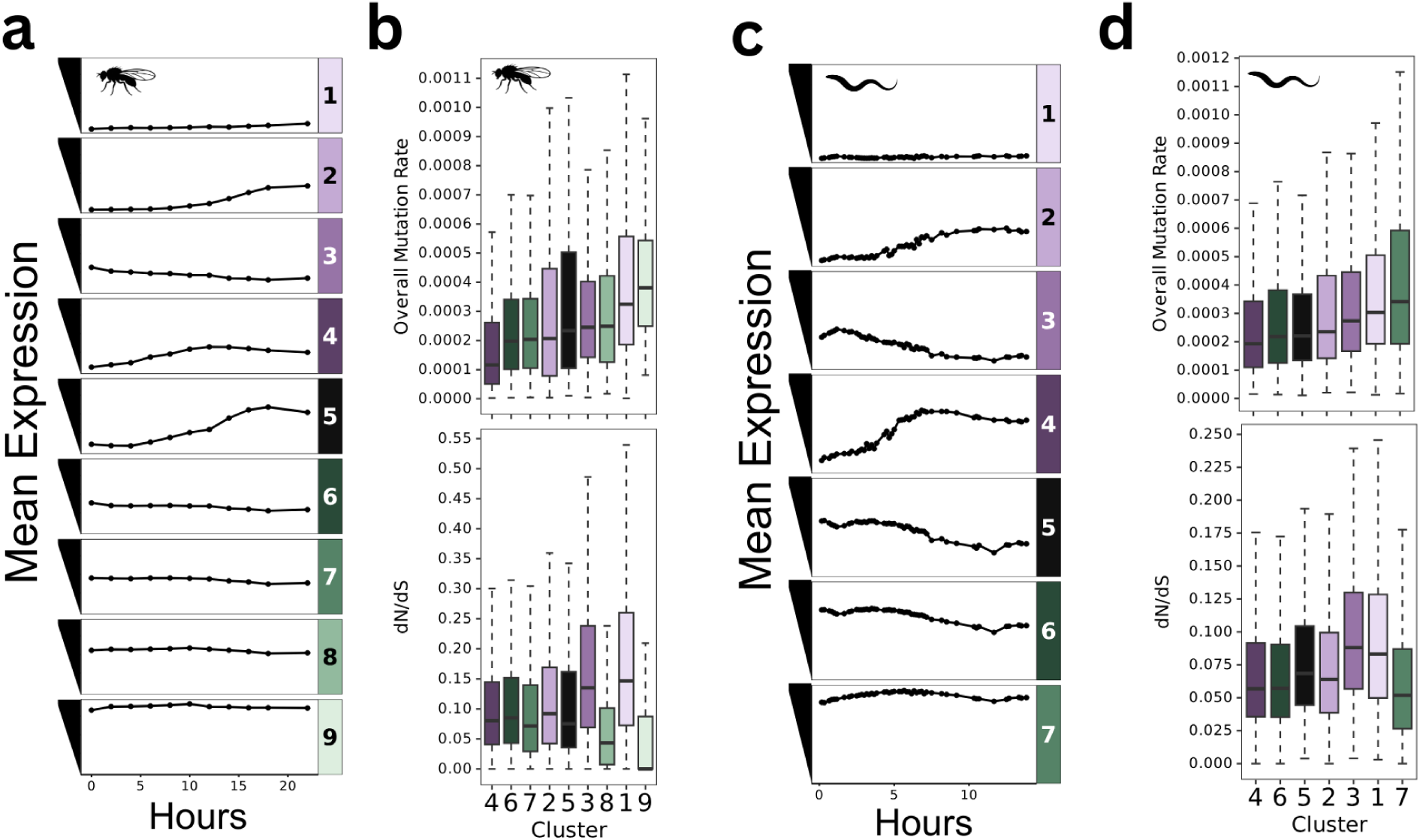
Distinct developmental expression programmes exhibit different mutation rates. **(a,c)** Line plots showing mean expression trajectories of *k*-means clusters of developmental gene expression in *D. melanogaster* **(a)** and *C. elegans* **(c)**. **(b,d)** Box plots showing matched all-type mutation rates (top) and dN/dS (bottom) of genes within each developmental expression cluster in *D. melanogaster* **(b)** and *C. elegans* **(d)**. Box plots show the median and interquartile range, with whiskers extending to 1.5× the interquartile range.

### 5.3 Embryonic gene expression is a stronger predictor of mutation rates than chromatin modifications

Previous studies in plants (**Monroe et al., 2022**) reported that genes under the strongest selective constraint and with the lowest mutation rates are enriched for specific chromatin modifications associated with active transcription, particularly H3K36me3 and H3K4me1. This was subsequently proposed to reflect the association of H3K4me1 with mismatch repair (**Quiroz et al., 2024**), providing a potential mechanistic explanation for mutation bias. To test whether a similar mechanism might explain the association between mutation rate and expression in metazoans, we investigated whether chromatin state, together with other covariates including GC content, replication timing, and recombination rates were associated with variation in gene-body mutation rates, and compared these with gene expression.

Chromatin marks associated with repression were positively correlated with mutation rate, whereas activating marks were negatively correlated (**Fig. 4a–b, left panels**). Nevertheless, embryonic gene expression showed the strongest overall negative correlation with mutation rate, while GC content showed the strongest positive correlation (**Fig. 4a–b, left panels**). However, these predictors were themselves correlated, making it difficult to determine their independent contributions from individual correlations alone. We therefore employed LASSO regression to account for covariance among predictors. Developmental gene expression remained the strongest independent predictor of reduced mutation rate in both species (**Fig. 4a–b, right panels**), supporting developmental gene expression as a major predictor of genome-wide variation in mutation rate.

**Figure 4.**
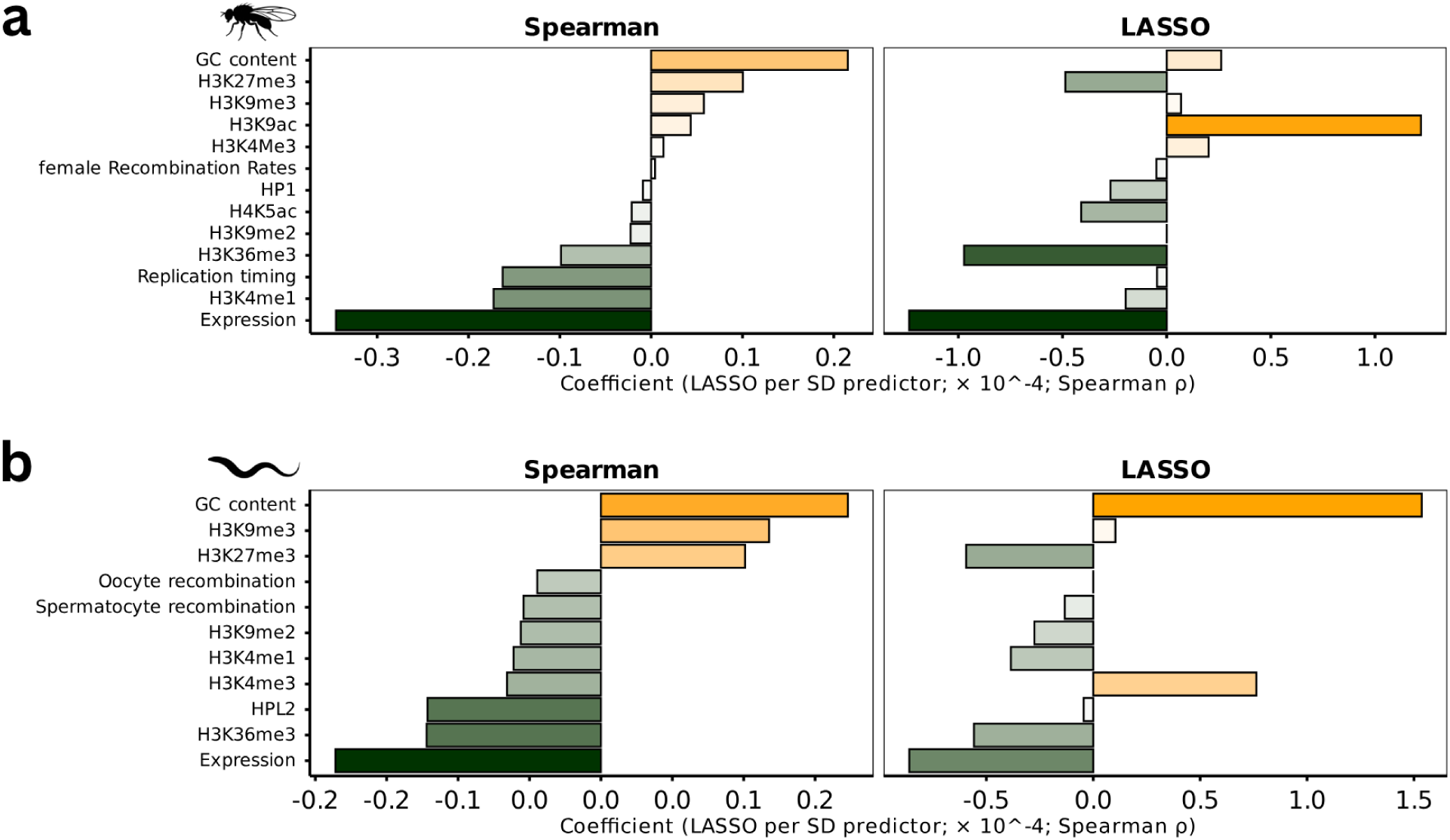
Gene expression explains mutation-rate variation better than epigenetic modifications. **(a,b)** Paired bar plots showing (left) univariate Spearman correlation coefficients between protein-coding gene mutation rates and mean gene-body epigenetic modification levels or whole-gene properties (gene expression and GC content), and (right) the standardized coefficients retained following LASSO regularization, for *D. melanogaster* **(a)** and *C. elegans* **(b)**.

### 5.4 Gene Mutation Rates Reflect Levels of Transcription-Coupled Repair

The above data strongly suggested that gene expression itself, rather than chromatin marks associated with transcription, was the most important factor predicting lower mutation rates. Whilst mismatch repair is linked to chromatin modification (**Makova and Hardison, 2015**), transcription is associated with increased rates of DNA repair due to the transcription coupled repair pathway (TCR). TCR is a sub-pathway of Nucleotide Excision repair involving the dedicated TCR proteins CSA and CSB which recruit the core excision repair complex (XP proteins) to stalled RNA polymerase (**Selby et al., 2023)**. The global genome-wide NER pathway, in contrast, is independent of polymerase stalling as it detects damage directly via the protein XPC (**Nieto Moreno, Olthof and Svejstrup, 2023**). To test whether TCR might be involved in reducing mutation rate at transcribed genes, we took advantage of XR-seq data for *D. melanogaster* (**Deger *et al*., 2022**) and *C. elegans* (**Kose *et al*., 2025**). XR-seq sequences the excised fragment after it is removed by the NER complex, thus an increased signal reflects increased ongoing repair by either TCR or global genome-wide NER (**Hu et al., 2019**).

In both species, excision repair was enriched across gene bodies, with the greatest repair signals being found in the most highly expressed genes (**Fig. S10a-b**) and genes with the lowest mutation rates (**Fig. 5a-b**). This could reflect either TCR or global genome repair. XPC is a component specific for global genome repair, thus knockouts of XPC should only undergo TCR. In *C. elegans* XPC knockout animals, where only transcription-coupled repair remains active, excision repair was still associated with expression (**Fig. S10c**) and mutation rate (**Fig. S10d**). Conversely, in CSA knockout worms too, where only global XR is active, excision repair was still associated with expression (**Fig. S10e**) and mutation rate (**Fig. S10f**). However, in *D. melanogaster* XPC knockout animals, excision repair no longer tracked with expression level (**Fig. S10g**), but still tracked with mutation rate (**Fig. S9h**). Some studies in *Drosophila* have shown that XPC is involved in TCR as well as global genome repair (**Deger et al., 2022**), confounding the interpretation of these findings. Both *C. elegans* and *D. melanogaster* thus show strong evidence that NER is associated with genes with higher developmental expression and this may explain their reduced mutation rate; at least in *C. elegans* this likely reflects direct recruitment of DNA repair by stalled RNA polymerase in TCR; however, the persistence of the relationship in the absence of TCR suggests that NER itself may also be involved in promoting repair of highly transcribed genes.

**Figure 5.**
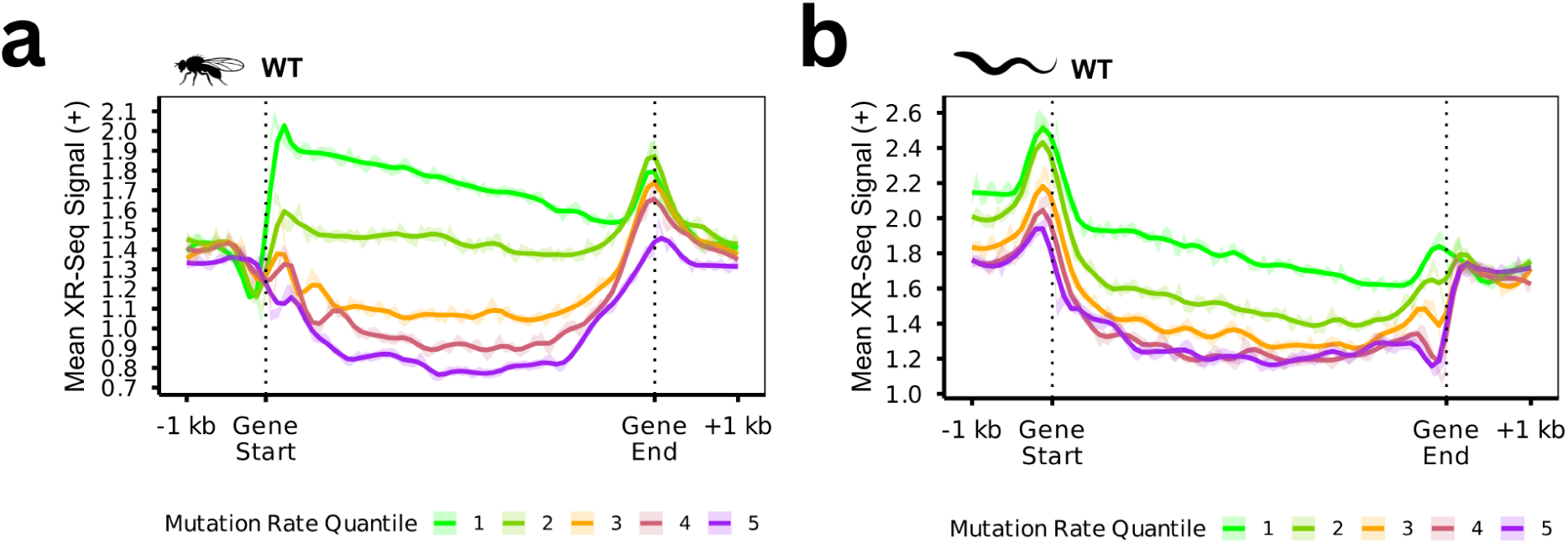
Reduced mutation rates in highly expressed genes are associated with elevated nucleotide excision repair. Relative mean nucleotide excision repair (NER) profiles across gene bodies for genes grouped into mutation-rate quintiles (1 = lowest mutation rate, 5 = highest mutation rate) in **(a)** wild-type *D. melanogaster* and **(c)** wild-type *C. elegans*. Dashed lines indicate gene start and end sites.

### 5.5 Mutation Rate Divergence Reflects Embryonic Gene Expression Divergence

These data suggest that developmental gene expression was the most strongly associatedof factor with genome-wide mutation-rate variation, most likely due to increased nucleotide excision repair, including TCR. This hypothesis would predict that changing developmental gene expression would affect mutation rate. To test this we took advantage of evolution. We examined MA data from the nematode *P. pacificus* (see Methods), a relative of *C. elegans* which diverged from *C. elegans* approximately 100Mya (**Weller et al., 2014**; **Wang et al., 2026**). *P. pacificus* shares highly similar developmental gene expression profiles and chromatin landscape with *C. elegans* (**Werner et al., 2018; Beltran et al., 2019**).

We first verified that the main patterns observed in *C. elegans* and *D. melanogaster* were also present in *P. pacificus*: genes under stronger purifying selection had lower mutation rates (**Fig. S10a**), embryonic expression was negatively associated with mutation rate across development (**Fig. S10b**), and expression remained the strongest predictor after accounting for other epigenetic features (**Fig. S10c**). We thus predicted that genes that change in their developmental profile between *C. elegans* and *P. pacificus* might show corresponding changes in mutation rate. To test this we obtained 1:1 orthologues between *C. elegans* and *P. pacificus*. Note that the mutation rate estimations are not absolute mutation rates due to the differing numbers of generations and number of lines in the studies, but we could compare how the relative mutation rate differed compared to the baseline mutation rate between species.

Changes in relative mutation rate were strongly negatively correlated **(ρ = -0.12, P = P < 4.39 x 10^-8^)** with changes in relative expression (**Fig. S12a**) across the entire distribution of relative expression changes, as revealed by binning orthologs into equal-sized quantiles based on their expression change (**Jonckheere P < 0.001; Fig. 6a**). To test whether there might still be a stronger association between either up or down-regulation amongst orthologs that diverged the most, we compared the change in relative mutation rate of orthologs which had either very large increases (>60%) or decreases (<60%) in expression against a down sampled subset of orthologs with negligible (<20%) change in expression (**Fig. 6b**). Both directions showed significant differences, however increases in expression resulted in more significant decreases in mutation rate (**Wilcoxon: *p* = 0.009**) than decreasing expression on increasing mutation rates (**Wilcoxon: *p* = 0.021**). We noted that even orthologues with negligible expression change (<20% percentile change) exhibited higher mutation rates in *P. pacificus* than in *C. elegans*, due to a higher baseline rate (see above). To circumvent this, we tracked expression changes across within-species expression quantiles re-calculated using only orthologs with expression in both species. Dropping expression quantile resulted in a statistically significant increase in mutation rate (**Wilcoxon: P < 0.001**) whereas increasing expression quantile was not associated with mutation rate (**Wilcoxon: P = 0.10; Fig. 6c**). Moreover, increasingly large downward steps were associated with progressively larger increases in relative mutation rate, and vice versa (**Jonckheere P = 2.86 x 10^-6^; Fig. 6d**). Importantly, changes in mutation rate associated with transcription were not matched by equivalent changes in dN/dS **(Fig. S12b**), either across the entire binned distribution (**Fig. S12c**), only at the extremes (**Fig. S12d**), or following expression quantile-changes (**Fig. S12e-f)**. This argues that transcriptional changes across evolution may lead to differences in mutation rate, and that this can occur independently of changes in selective constraint.

**Figure 6.**
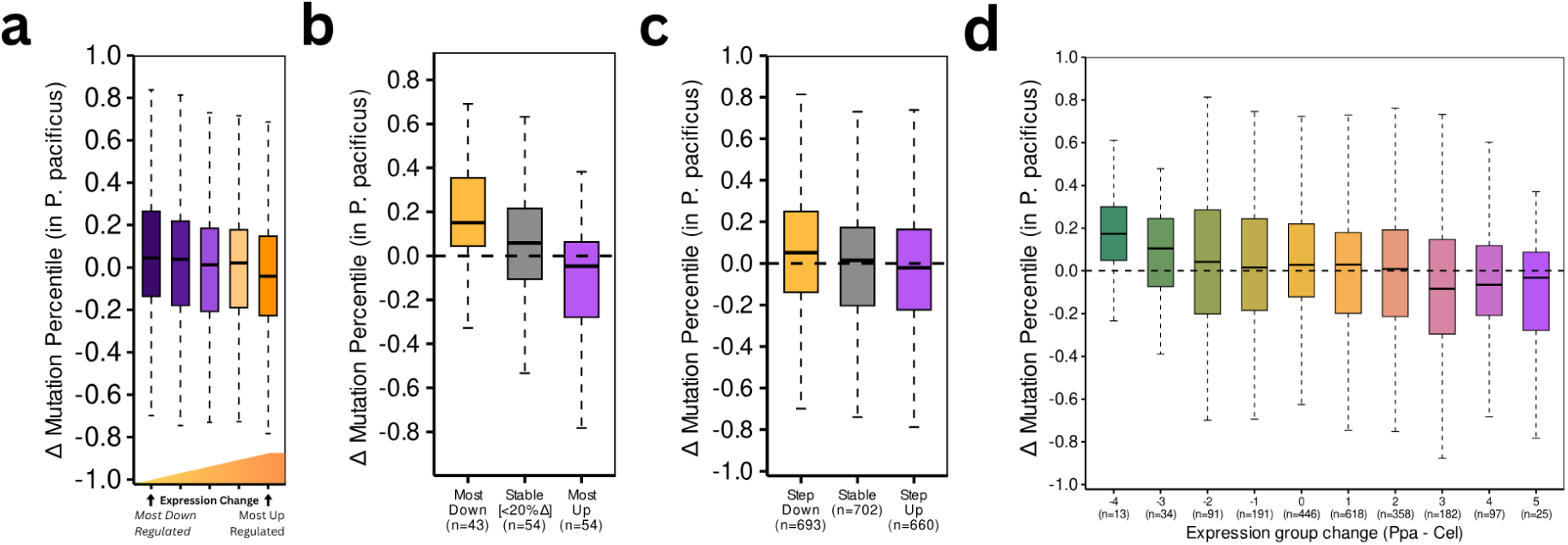
Evolutionary divergence in embryonic gene expression predicts divergence in local mutation rates. **(a)** Relative mutation-rate divergence among orthologues grouped into quintiles of embryonic expression divergence in *Pristionchus pacificus* relative to *Caenorhabditis elegans*. **(b)** Relative mutation-rate divergence of orthologues exhibiting the largest (>60%) decreases (orange) or increases (purple) in embryonic expression compared with random down-sample of orthologues showing negligible (<20%) expression divergence (grey). **(c)** Change in relative mutation rate of orthologs decreasing in relative embryonic expression quantile (orange, Step Down), remaining in the same quantile (grey, Stable) or increasing in relative embryonic expression quantile (purple, Step Up). **(d)** Relative mutation-rate divergence as a function of the magnitude of embryonic expression-quantile change.

## Discussion

Genetic mutations are the primary source of phenotypic variability, and thus limit the pace and direction of adaptation. Traditionally, mutations have been considered to occur at random with respect to their effect on fitness. However, advances in genomic sequencing technologies have provided unprecedented characterisation of genetic variability and reveal that loci under the strongest selective constraint may accumulate mutations more slowly, whereas loci under diversifying selection accumulate more mutations. Here we produce further evidence for such germline “adaptive mutation bias”, extending observations previously reported in bacteria (**Martincorena, Seshasayee and Luscombe, 2012)** and plants (**Monroe et al., 2022**) to two distantly related animal species, *D. melanogaster* and *C. elegans*. We utilised MA experiment data to directly assay both coding and putatively neutral DNVs, largely bypassing the historical selection that confounds estimates of DNVs in natural populations that has severely limited most previous investigations.

Previous work in *Arabidopsis* proposed that chromatin modifications, specifically H3K4me1, reduce local mutation rates via recruitment of mismatch repair proteins (**Monroe et al., 2022; Quiroz et al., 2024**). Although several chromatin features were indeed associated with local mutation rates in *D. melanogaster* and *C. elegans*, including H3K4me1, we found that developmental gene expression instead explained substantially more variation than any single chromatin modification, even after accounting for covariance between epigenetic and transcriptional processes using LASSO regression. The XR-seq analyses provided a plausible mechanistic explanation for this relationship. Genes most depleted of mutations were consistently observed to have the greatest nucleotide excision repair profiles. In *C. elegans*, the fact that this was observed strongly in XPC knockout animals, in which the only form of NER is TCR, suggests that TCR could explain the preferential repair, and thus reduced mutation rate, at these loci. It is important to note that in CBS knockout *C. elegans*, where only global XR is active, the relationship between XR-seq signal and both expression and mutation rate were still present, although diminished in strength. Thus our data might suggest an additional preference for global genome repair to be associated with transcription. Increased NER has been associated with open chromatin in analyses of mutation rates in human cancer (**Polak et al., 2014**), which might provide an additional mechanism preferentially targeting excision repair to highly transcribed genes. Conversely, in XPC knockout *D. melanogaster*, XR-seq signal no longer reflected expression, indicating loss of TCR consistent with TCR being dependent on XPC in *D. melanogaster* (**Deger et al., 2022**). Despite this, XR-seq signal was still greatest at loci with the lowest mutation rates in XPC knockout *D. melanogaster*, consistent the existence of an XPC-independent XR pathway, although alternative explanations involving confounding genomic features and non-specific XR-seq signal remain to be excluded.

An additional qualification worth noting is that mutation rate did not reduce monotonically as expression increased, but instead began to increase amongst the most highly expressed loci. Indeed, loci which were consistently extremely highly expressed across all of development exhibited nominally higher mutation rates than even the most lowly expressed loci. This may be consistent with previous reports of a trade-off between TCR and transcriptional-coupled damage (TCD; **Xia et al., 2020**). Taken together, therefore, we propose that reduced mutation rate at genes that are under purifying selection in *C. elegans* and *Drosophila* is due to the developmental expression profile of these genes, which causes them to be preferentially targeted by excision repair processes.

Our results are important to consider in the light of the debates over whether mutation bias can ever be the direct product of evolution by natural selection (see for example **Lynch et al., 2023**). The existence of a mechanism to explain the observed mutation bias might suggest that the process has evolved. Theoretical descriptions consider ‘trans-acting modifiers’: alleles which act to reduce mutation rate at a selection of genes, and that these alleles may go to fixation in the population due to the reduced mutational burden that accumulates. In one interpretation of our data, the association of NER with active transcription, most obviously through TCR but potentially through global NER as well, would itself be considered such a trans-acting modifier. However, it is important to consider that the beneficial effect associated with increased repair at transcribed regions may not be limited to prospective reductions in mutation load. Mutations in early embryogenesis in important developmental genes, such as premature stop codons, might lead directly to developmental abnormalities, independent of any lasting effect on mutational burden. Moreover, the presence of certain types of damage in the transcribed regions may itself negatively impact transcriptional rates, again leading to developmental abnormalities. The fact that our data suggests that the relationship between germline mutation rate and transcription applies to genes that are expressed in other embryonic lineages rather than the germline strongly suggests that other selective advantages beyond mutation load may be at play. Thus it is possible that the mutation bias that we observe, including the relationship between purifying selection and low mutation rate, may be a consequence of selection for developmental robustness, potentially through avoiding somatic mutations early in development in highly expressed genes, rather than direct selection on the mutation rate. An important point in support of this argument is the fact that expression divergence between *C. elegans* and *P. pacificus* was clearly associated with mutation rate changes, but was not clearly associated with changes in dN/dS, which shows that, at least over relatively short evolutionary timescales, mutation rate shifts can be decoupled from selective constraint. Nevertheless, it is intriguing that the same type of mutation bias (lower mutation rate at genes with housekeeping functions) is seen in both plants and animals, but with different mechanisms responsible. Arguably, this strengthens the possibility that this form of mutation bias is indeed shaped by evolution - it has evolved convergently in distinct lineages. Rigorous analyses of mutation bias by studying mutations in the absence of selection in other eukaryotic taxa will be important to test this conclusion.

A number of limitations of our work need to be considered. Principally, MA experiments minimise but do not eliminate natural selection. It is therefore possible that DNV rate estimates are at least partially shaped by selection. Indeed, in a recent MA study using *Daphnia*, mutation bias was identified, but not at synonymous sites, suggesting that selection can influence mutation bias even in MA experiments with reduced population size (**Deng et al., 2026**). Our data showed mutation bias even in synonymous sites. Nevertheless, even intronic (**Girardini, Olthof and Kanadia, 2023**) and synonymous (**Zhang and Qian, 2025**) sites are not entirely isolated from fitness. The only way to fully address this is to establish mutation rates in single individual germlines, something which currently remains impossible but, with new sequencing technologies with the potential to assess mutations in individual strands of DNA, could become accessible in the future.

An important additional limitation is our use of *P. pacificus* and *C. elegans* to demonstrate how changes in transcription over evolution led to changes in mutation rate. Whilst these species are relatively close, they are still separated by 100my and as a result relatively few orthologous genes could be compared. Although MA lines have been constructed from closer relatives to *C. elegans* (**Chen et al., 2023**), unfortunately we were not able to use these due to paucity of mutations identified in these lines (100s rather than 1000s).

Our analysis was restricted to comparative study; however, mechanistic confirmation of the importance of NER would be a next step for this research program. In particular, long term MA lines containing thousands of mutations made in NER mutant lines would be an advantage; additionally, combining NER mutants with single molecule assessment of mutation rate (see above) would be a good way to test our mechanistic model.

In summary, our findings reinforce the view that mutation rates are non-random with respect to their fitness consequences and propose strong evidence that their evolution is shaped more by local transcriptional regulation, rather than local epigenetic environment. Adaptive germline mutation bias may therefore emerge indirectly as a consequence of developmental gene expression evolution, providing a mechanistic framework that reconciles adaptive mutation bias with theoretical limits on the direct evolution of mutation rates and thus greatly increasing its phylogenetic generality.

## Acknowledgements

We thank Tobias Warnecke and Lars Jansen for their feedback and constructive criticism, as well as Denis Noble and all members of the Sarkies EpiEvo Group for many engaging discussions and suggestions. We also thank Natalie Jones for assisting in proofreading and formatting the manuscript.

## Data availability

All data used as part of this project were acquired from previously published studies or publicly available repositories. Accession numbers and sources for mutation accumulation datasets, genome annotations, gene expression datasets, epigenomic datasets and other genomic covariates are provided in the Methods and Supplementary Table 1. Processed data underlying the analyses and figures in this study, including gene-level mutation-rate estimates, orthology and processed expression and genomic-feature datasets, as well as analysis pipeline and scripts required to reproduce the results and figures are available at [https://github.com/danphillips28/Panspecies_MA_Analyses]. No new sequencing data were generated as part of this study.

## Supplementary Materials

**Figure S1.**
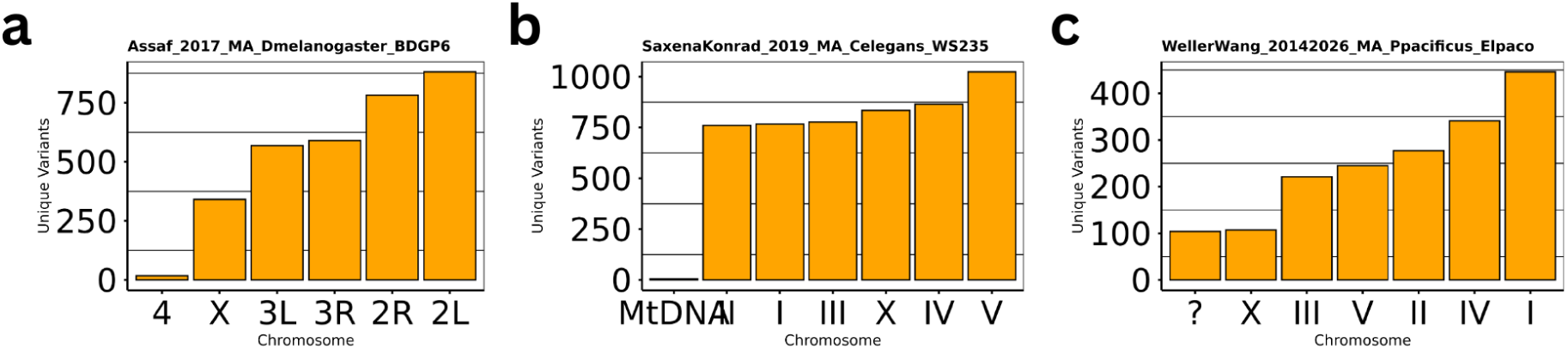
Unique *de novo* variants identified on each chromosome in **(a)** *D. melanogaster* **(b)** *C. elegans* and **(c)** *P. pacificus* following mutation accumulation experiments.

**Figure S2.**
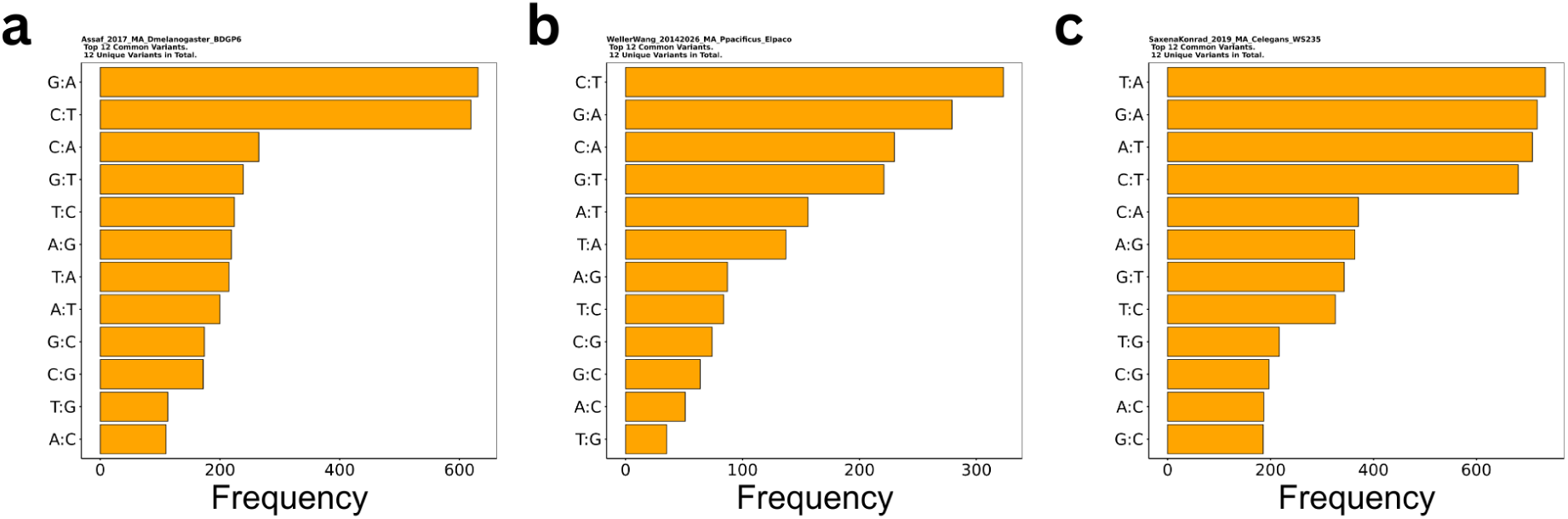
Unique de novo variants of different substitution types identified following mutation accumulation experiments in **(a)** *D. melanogaster* **(b)** *P. pacificus* and **(c)** *C. elegans*.

**Figure S3.**
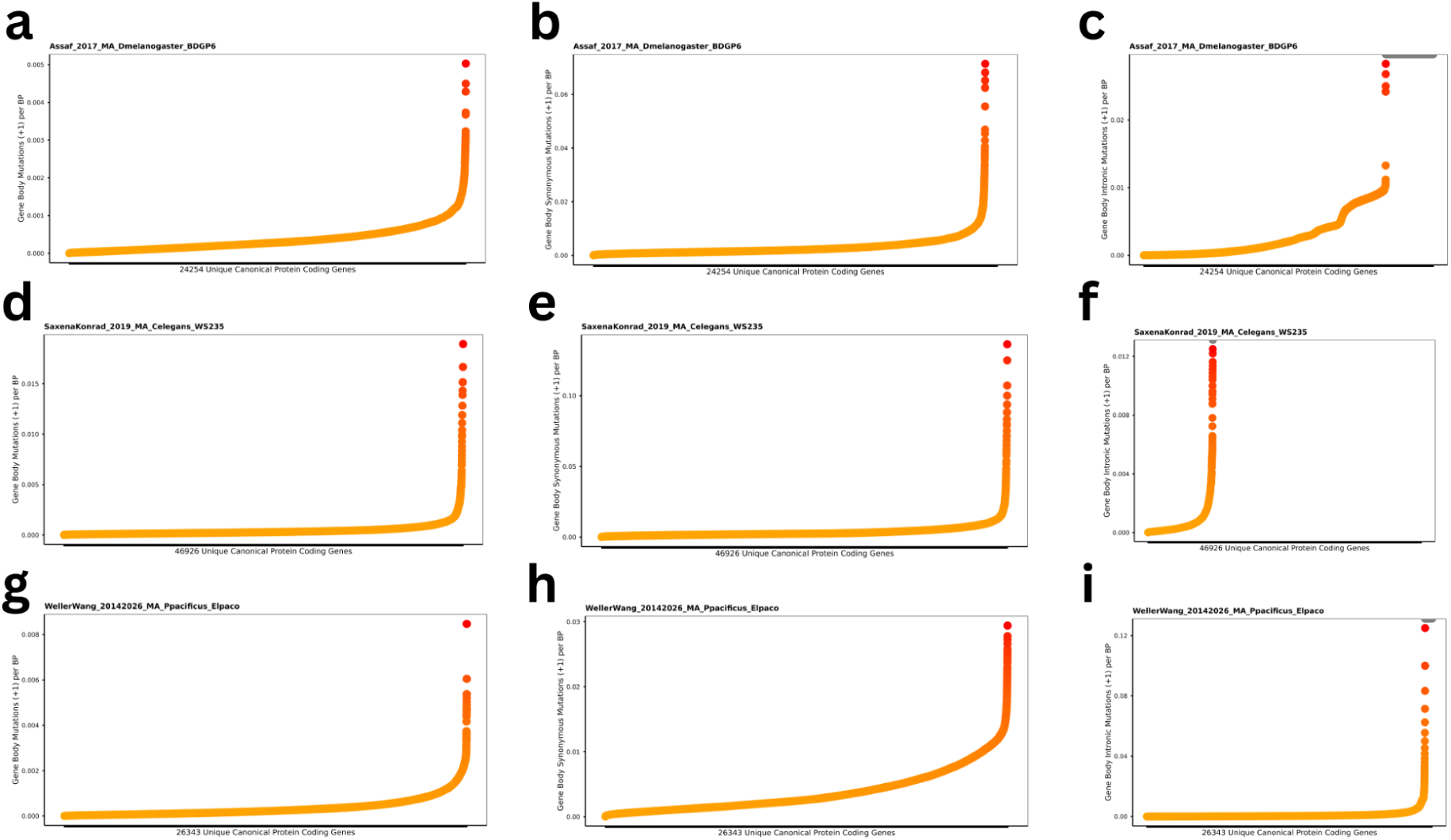
Scatter plots showing the all-type **(a,d,g)**, synonymous **(b,e,h)** and intronic **(c,f,i)** mutation rate distribution of protein coding genes in **(a,b,c)** *D. melanogaster* **(d,e,f)** *C. elegans* and **(g,h,i)** *P. pacificus*.

**Figure S4.**
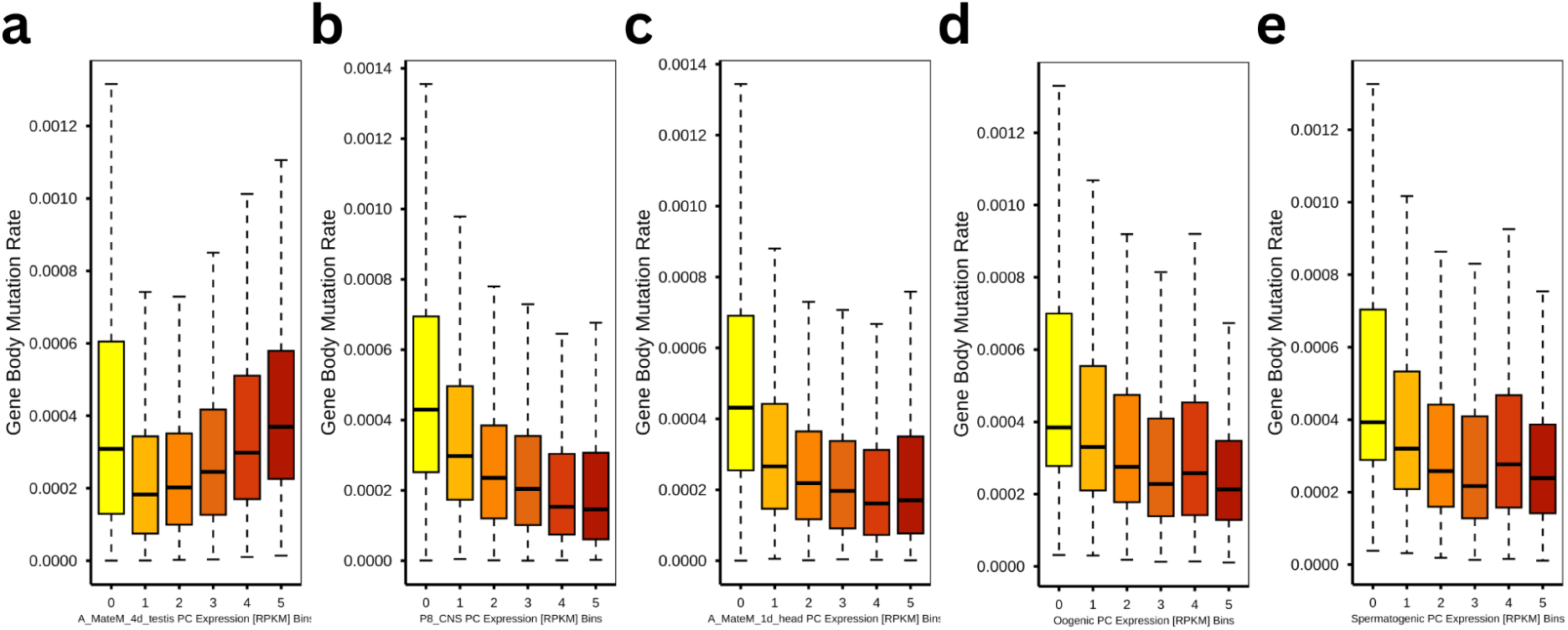
Mutation rates of genes grouped into expression quintiles from **(a)** *D. melanogaster testis* **(b)** *D. melanogaster* central nervous system **(c)** *D. melanogaster* head **(d)** *C. elegans* oogenic gametes and **(e)** *C. elegans* spermatogenic gametes. Box plots show the median and interquartile range, with whiskers extending to 1.5× the interquartile range.

**Figure S5.**
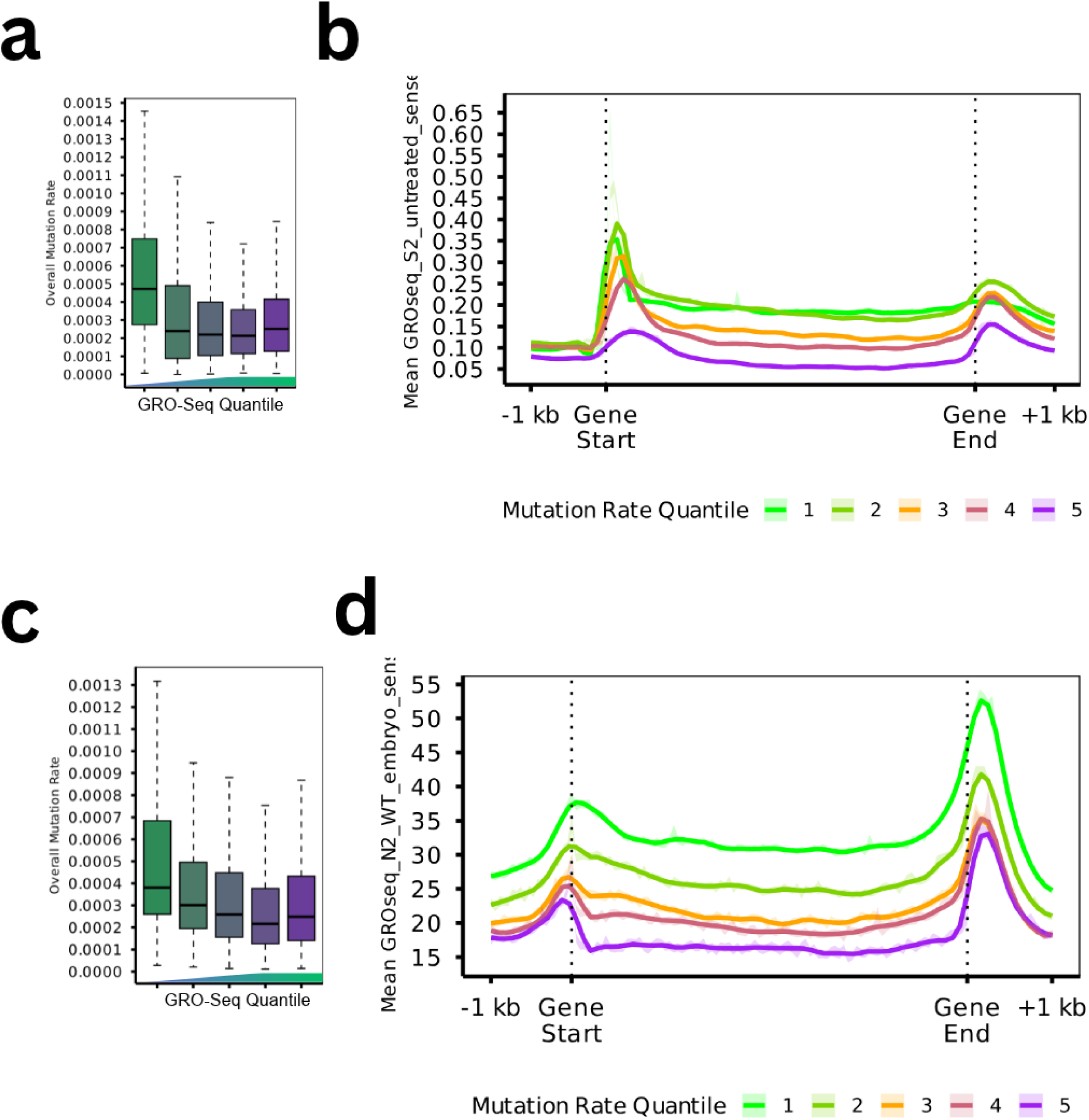
Mutation rates of genes grouped into Global Run-On sequencing (GRO-Seq) signal quantiles from **(a)** *D. melanogaster* S2 cells and **(c)** *C. elegans* early embryos. Box plots show the median and interquartile range, with whiskers extending to 1.5× the interquartile range. Relative mean GRO-Seq profiles across gene bodies for genes grouped into mutation rate quantiles in **(b)** *D. melanogaster* S2 cells and **(d)** *C. elegans* early embryos. Mutation-rate quantiles (1 = lowest mutation rate, 5 = highest mutation rate).

**Figure S6.**
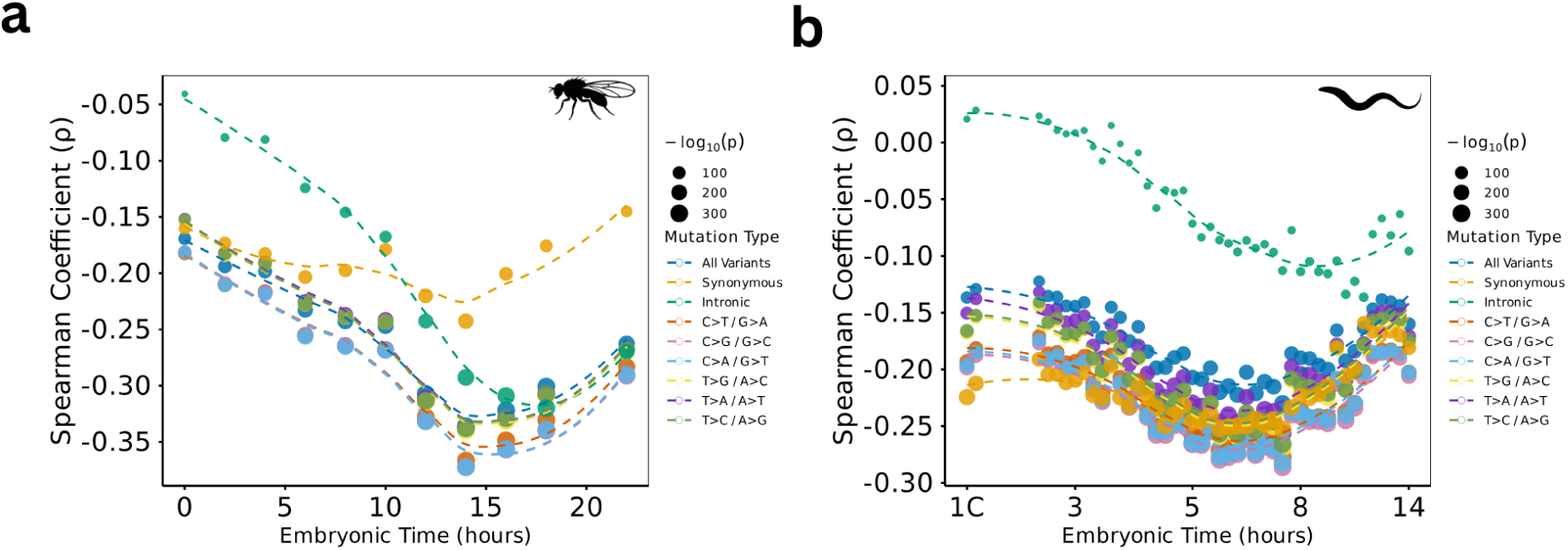
Line plots showing Spearman correlation coefficients between protein-coding gene expression and mutation rate across developmental time in *D. melanogaster* **(a)** and *C. elegans* **(b)**. Colours indicate different mutation categories (all mutations, synonymous mutations, intronic mutations, and the six substitution classes), while point size represents −log^10^(adjusted P value).

**Figure S7.**
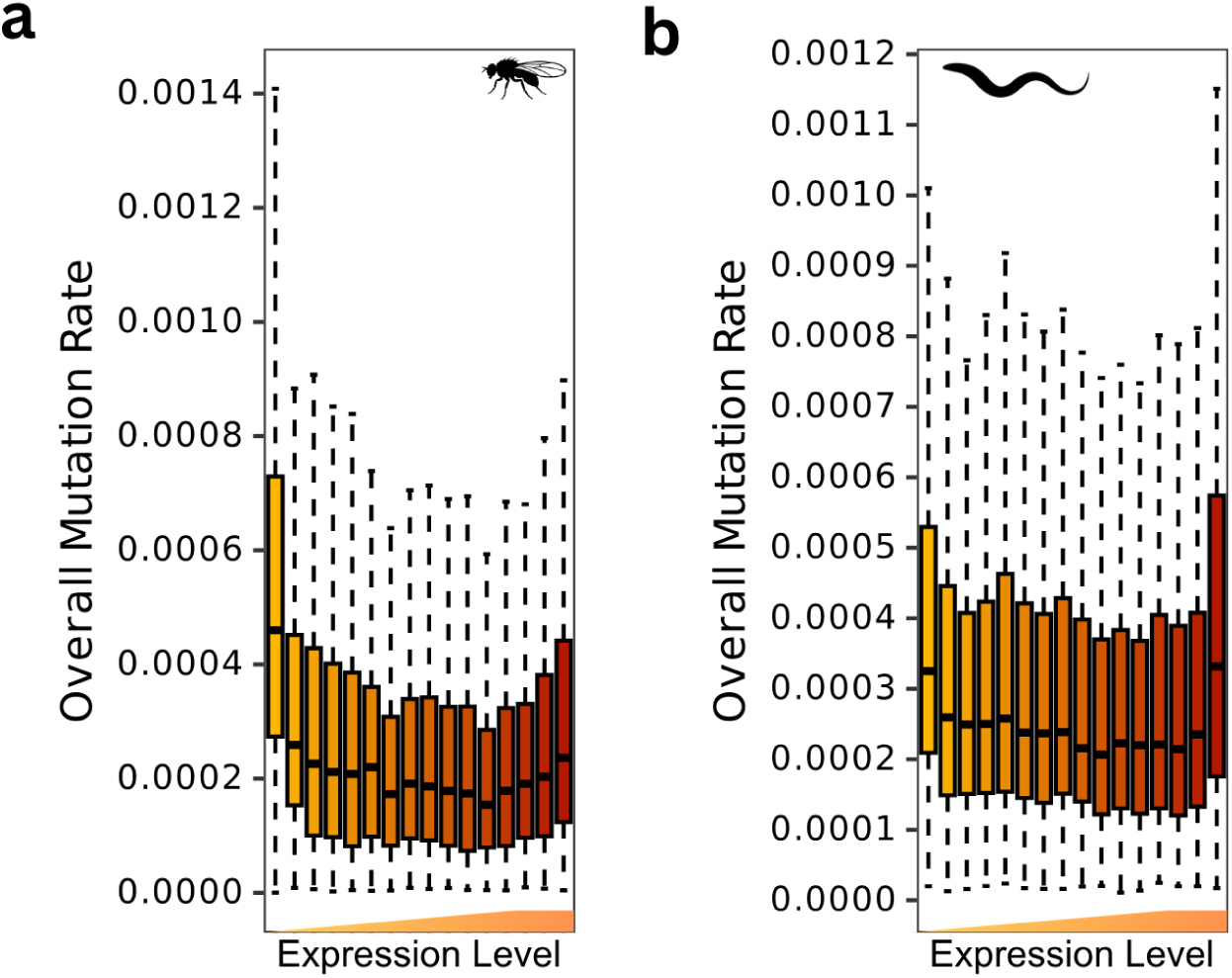
Bar plots showing mutation rates of genes grouped into 16 expression quantiles according to late (14-16hr) developmental expression in *D. melanogaster* **(a)** or mid (430min) developmental expression in *C. elegans* **(b)**. Box plots show the median and interquartile range, with whiskers extending to 1.5× the interquartile range.

**Figure S8.**
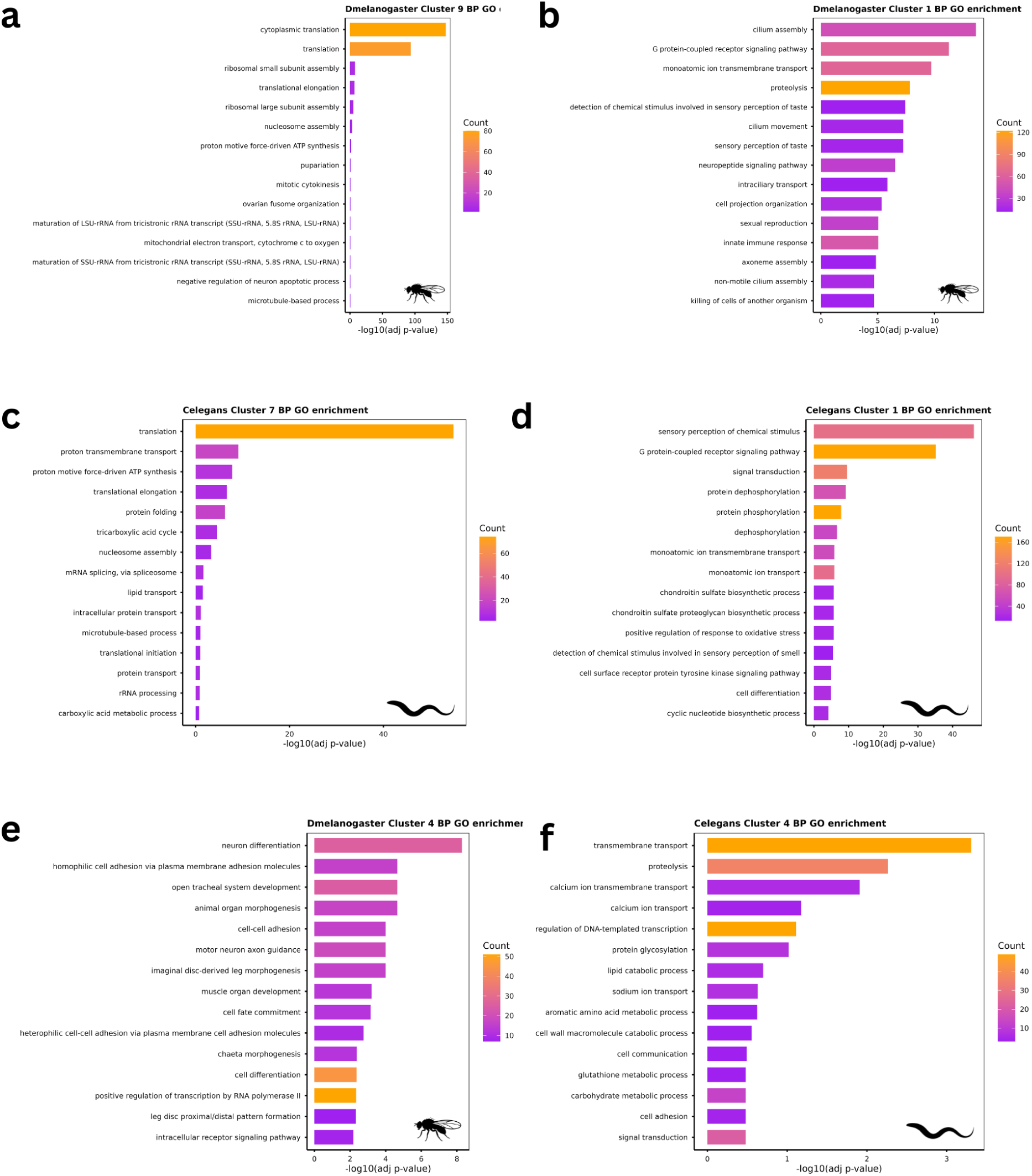
Bar plots showing the over-representation of biological process (BP) terms in **(a)** *D. melanogaster* K-means cluster 9 (high expression-high mutation rate) **(b)** *D. melanogaster* K-means cluster 1 (virtually no expression-high mutation rate) **(c)** *C. elegans* K-means cluster 7 (high expression-high mutation rate) **(d)** *C. elegans* K-means cluster 1 (virtually no expression-high mutation rate) **(e)** *D. melanogaster* K-means cluster 4 (mid-developmental expression peak-lowest mutation rate) and **(f)** *C. elegans* K-means cluster 4 (mid-developmental expression peak-lowest mutation rate).

**Figure S9.**
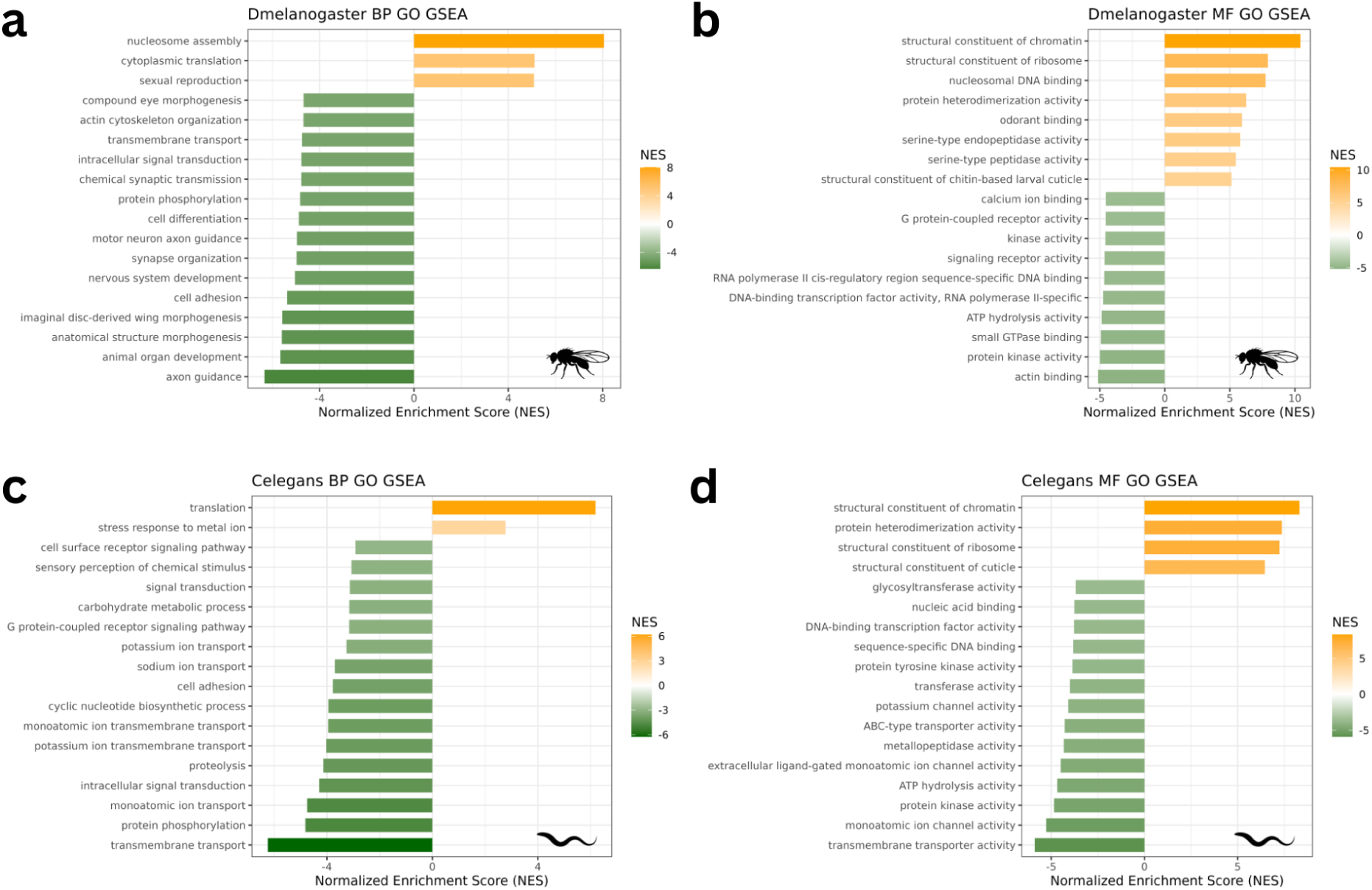
Bar plots showing association between mutation rates and **(a)** biological processes in *D. melanogaster* **(b)** molecular functions in *D. melanogaster* **(c)** biological processes in *C. elegans* **(d)** molecular functions in *C. elegans* identified using gene set enrichment analysis (GSEA) of mutation rate distributions calculated from mutation accumulation experiments.

**Figure S10.**
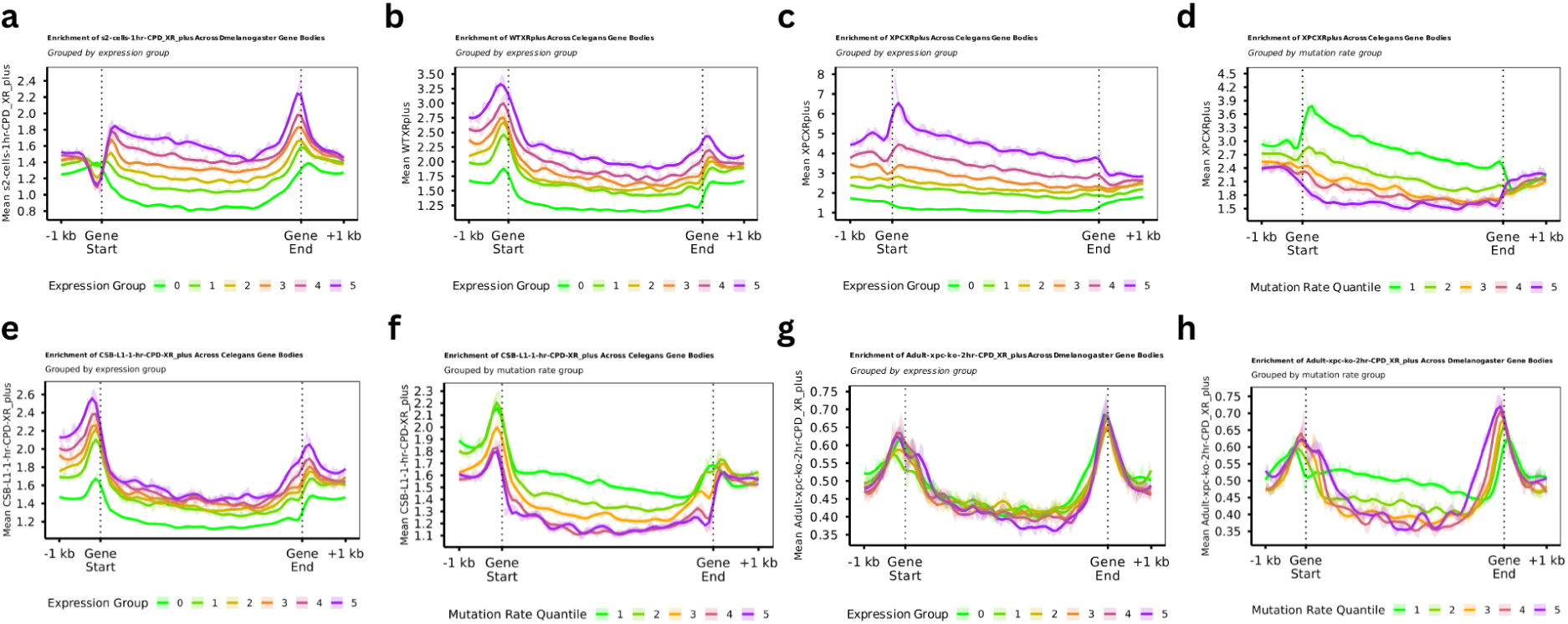
Relative mean nucleotide excision repair (NER) profiles across gene bodies for genes grouped into **(a)** expression quantiles in WT *D. melanogaster* **(b)** expression quantiles in *C. elegans* **(c)** expression quantiles in XPC-KO *C. elegans* **(d)** mutation rate quantiles in XPC-KO *C. elegans* **(e)** expression quantiles in CBS-KO *C. elegans* **(f)** mutation rate quantiles in CSB-KO *C. elegans* **(g)** expression quantiles in XPC-KO *D. melanogaster* **(h)** mutation rate quantiles in XPC-KO *D. melanogaster.* Mutation-rate quantiles (1 = lowest mutation rate, 5 = highest mutation rate) and expression quantiles (0 = no expression, followed by 5 quantiles of increasing expression).

**Figure S11.**
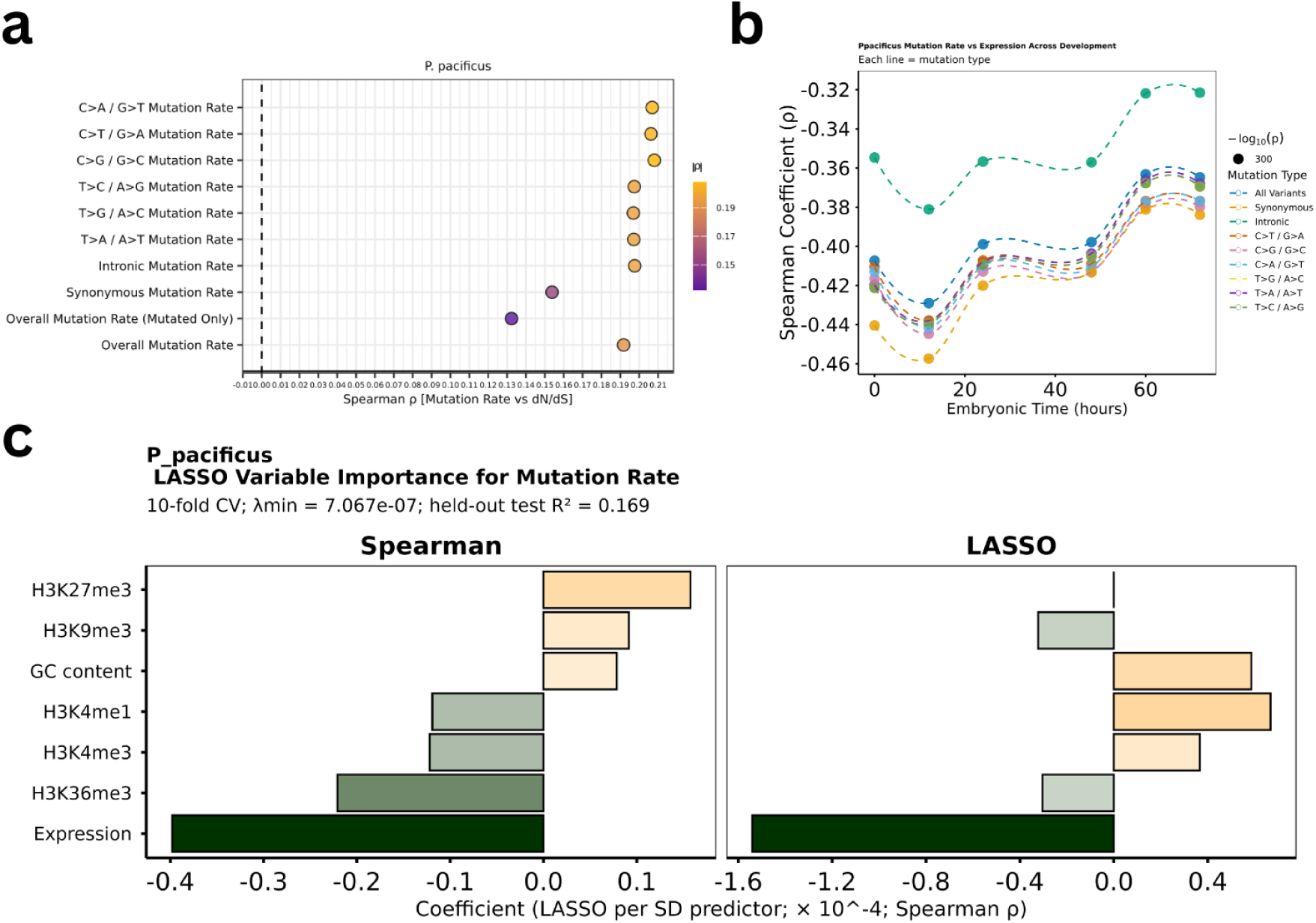
**(a)** Spearman correlation coefficients between log(mutation rate) and log(dN/dS + 0.5) for each mutation class in *P. pacificus* following mutation accumulation experiment.. Point size represents –log₁₀(adjusted P value), and colour indicates correlation coefficient. (b) Line plots showing Spearman correlation coefficients between protein-coding gene expression and mutation rate across developmental time in *P. pacificus*. Colours indicate different mutation categories (all mutations, synonymous mutations, intronic mutations, and the six substitution classes), while point size represents −log10(adjusted P value). (c) Paired bar plots showing (left) univariate Spearman correlation coefficients between protein-coding gene mutation rates and mean gene-body epigenetic modification levels or whole-gene properties (gene expression and GC content), and (right) the standardized coefficients retained following LASSO regularization in *P. pacificus*.

**Figure S12.**
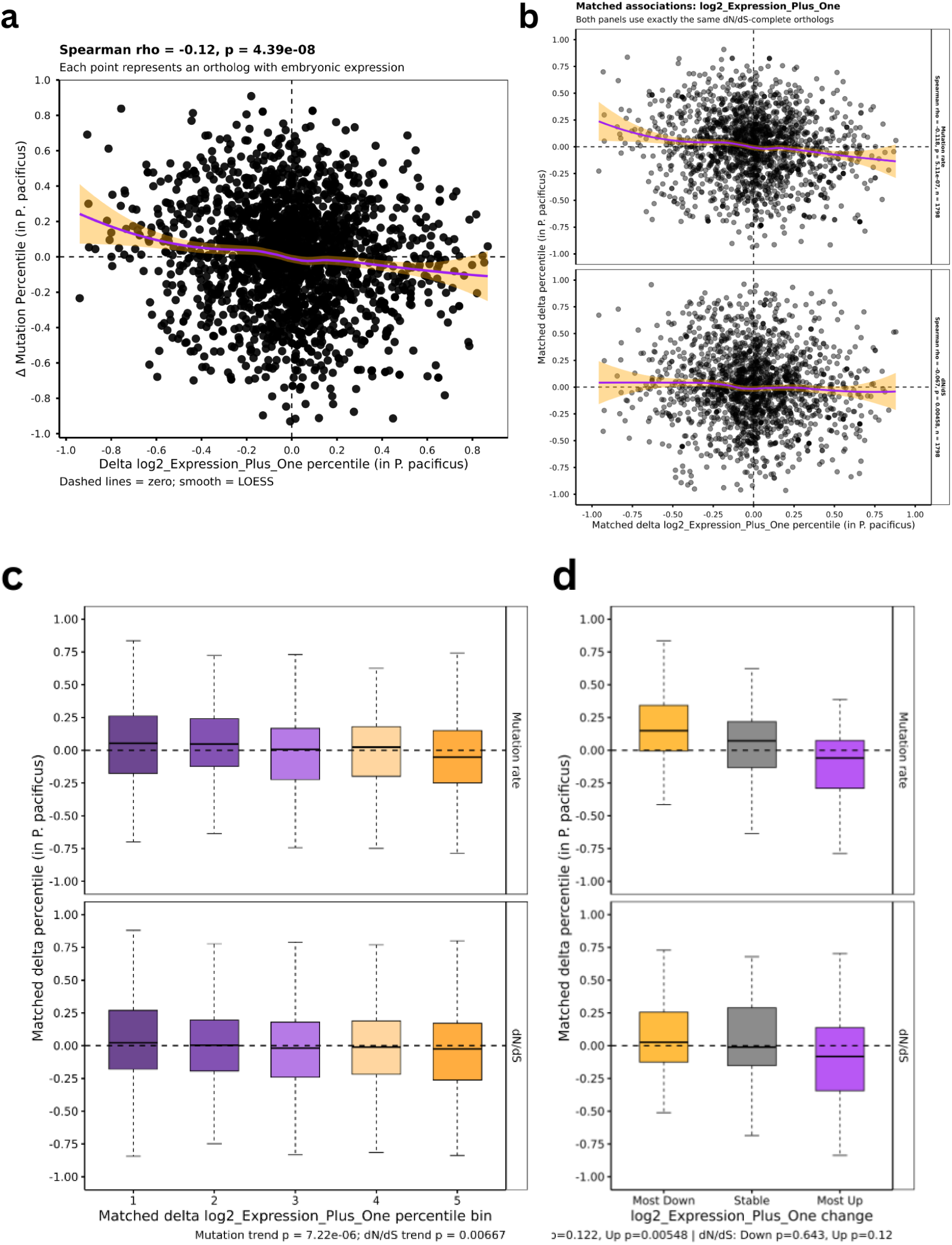
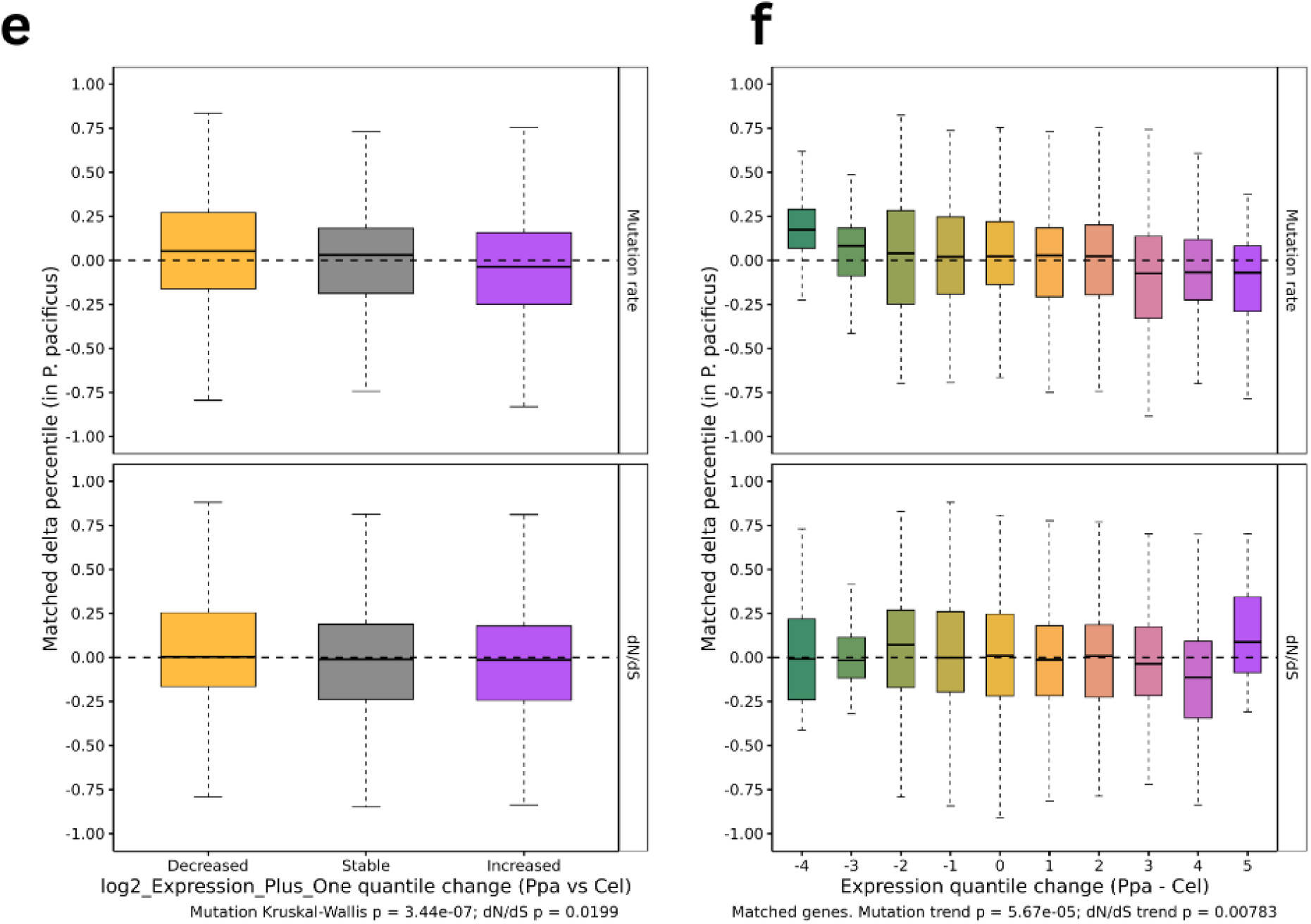
**(a)** Scatter plot showing the change in embryonic expression between orthologs shared between *P. pacificus* and *C. elegans* and their change in mutation rate. **(b) (top)** Same as **(a)** but restricted to orthologs with dN/dS estimates. **(bottom)** Scatter plot showing change in dN/dS of orthologs (compared to a close relative of each species) between *P. pacificus* and *C. elegans* and their change in embryonic expression. **(c)** Boxplots showing relative **(top)** mutation-rate and **(bottom)** dN/dS divergence among dN/dS-matched orthologues grouped into quintiles of embryonic expression divergence in *P. pacificus* relative to *C. elegans*. **(d)** Box plots showing relative **(top)** mutation-rate and **(bottom)** dN/dS divergence of matched-dN/dS orthologues exhibiting the largest (>60%) decreases (orange) or increases (purple) in embryonic expression compared with random down-sample of orthologues showing negligible (<20%) expression divergence (grey). **(e)** Box plots showing change in relative **(top)** mutation rate and **(bottom)** dN/dS of orthologs decreasing in relative embryonic expression quantile (orange, Step Down), remaining in the same quantile (grey, Stable) or increasing in relative embryonic expression quantile (purple, Step Up). **(f)** Relative **(top)** mutation-rate and **(bottom)** dN/dS divergence as a function of the magnitude of embryonic expression-quantile change.

**Supplementary Table 1.**

| **Species** | **Covariate** | **Source / accession** | **Original format** | **Genome build / conversion** | **LiftOver chain** | **Processing / replicates** |
|---|---|---|---|---|---|---|
| C. elegans | Expression | GSE50548 | .tab.gz | Gene IDs mapped using WormBase; WS235-labelled output |  | Embryonic_Expression generated from column 38 (430). |
| C. elegans | GC content | Calculated from reference sequence | Derived | WS235 |  | No external dataset accession required; calculation method to be documented in Methods. |
| C. elegans | GRO-seq (sense/antisense) | GSE43084; GSM1056279 (Kruesi et al., 2013) | .wig.gz → .bedgraph | WS230 (ce10 assumed) → WBcel235 (ce11) | ce10ToCe11.over.chain | N2 embryos; 1-bp plus/minus WIG tracks; 2 replicates pre-averaged and RPKM-normalised; WIG to bedGraph; sorted; no overlaps. |
| C. elegans | H3K27me3 | GSE114, GSE114440 | .bw | WS220 → WS235 | ce10ToCe11.over.chain.gz | Processed coverage track; coordinate conversion noted. Other processing not specified in supplied source table. |
| C. elegans | H3K36me3 | GSE114, GSE114440 | .bw | WS220 → WS235 | ce10ToCe11.over.chain.gz | Processed coverage track; coordinate conversion noted. Other processing not specified in supplied source table. |
| C. elegans | H3K4me1 | GSE114, GSE114440 | .bw | WS220 → WS235 | ce10ToCe11.over.chain.gz | Processed coverage track; coordinate conversion noted. Other processing not specified in supplied source table. |
| C. elegans | H3K4me3 | GSE114, GSE114440 | .bw | WS220 → WS235 | ce10ToCe11.over.chain.gz | Processed coverage track; coordinate conversion noted. Other processing not specified in supplied source table. |
| C. elegans | H3K9me2 | GSM2333097; GSM2333098 | .bw | ce10 (WS220) → ce11 (WS235) | ce10ToCe11.over.chain.gz | Mean of 2 replicates |
| C. elegans | H3K9me3 | GSM2515718; GSM2515719 | .bw | ce10 (WS220) → ce11 (WS235) | ce10ToCe11.over.chain.gz | Mean of 2 replicates |
| C. elegans | HPL2 | GSM2333101; GSM2333102 | .bw | ce10 (WS220) → ce11 (WS235) | ce10ToCe11.over.chain.gz | Mean of 2 replicates |
| C. elegans | Oocyte recombination | Bush et al. 2026; DOI 10.1371/journal.pgen.1012237, S1 | .csv → .bedgraph | GCA_028201515.1 → WS235 |  | minimap2 + paftools.js; non-overlapping 200-kb bins; cM/Mb |
| C. elegans | Replication timing | GSE140804 | .bed.gz | ce10 → ce11 | ce10ToCe11.over.chain.gz | Processed as Early – Late |
| C. elegans | Spermatocyte recombination | Bush et al. 2026; DOI 10.1371/journal.pgen.1012237, S1 | .csv → .bedgraph | GCA_028201515.1 → WS235 |  | minimap2 + paftools.js; non-overlapping 200-kb bins; cM/Mb |
| C. elegans | XR-seq (strand-specific) | GSM8595214/5215 (WT); GSM8595254/5255 (xpc-1); GSM8595234/5235 (CSB) | .bedgraph.gz → .bedgraph | Not specified in supplied script/manifest; no liftOver performed | Not applied in supplied script | L1 larvae, 1 h CPD; WT, xpc-1, CSB; 2 replicates per condition; compressed bedGraphs decompressed, sorted, averaged per strand with bedtools unionbedg (missing=0). |
| **Species** | **Covariate** | **Source / accession** | **Original format** | **Genome build / conversion** | **LiftOver chain** | **Processing / replicates** |
| D. melanogaster | Expression | FlyBase gene_rpkm_report_fb_2025_04 | .tsv.gz → .tsv | Gene-level IDs; no liftOver |  | Embryonic_Expression.tsv uses the 14–16 h embryonic time point. |
| D. melanogaster | Female recombination | Comeron Lab, R5/R6 crossover-rate workbook | .xlsx → .bedgraph | R6 / BDGP6 |  | R6 worksheet; non-overlapping intervals derived from 100-kb midpoints |
| D. melanogaster | GC content | Calculated from reference sequence | Derived | BDGP6 |  | No external dataset accession required; calculation method to be documented in Methods. |
| D. melanogaster | GRO-seq (sense/antisense) | GSE58955; GSM1423051; GSM1423052 (Fuda et al., 2015) | .bedgraph.gz → .bedgraph | dm3 (BDGP5) → dm6 (BDGP6) | dm3ToDm6.over.chain (where applied) | Untreated S2 cells; RPM-normalised; signed values converted to magnitudes before liftOver; 2 replicates averaged per strand (missing=0); sorted; no overlaps. |
| D. melanogaster | H3K27me3 | GSE20, GSE20781 | .bedGraph.gz | BDGP5 → BDGP6 | dm3ToDm6.over.chain.gz | Processed coverage track; coordinate conversion noted. Other processing not specified in supplied source table. |
| D. melanogaster | H3K36me3 | GSE20, GSE20785 | .bedGraph.gz | BDGP5 → BDGP6 | dm3ToDm6.over.chain.gz | Processed coverage track; coordinate conversion noted. Other processing not specified in supplied source table. |
| D. melanogaster | H3K4me1 | GSE20, GSE20786 | .bedGraph.gz | BDGP5 → BDGP6 | dm3ToDm6.over.chain.gz | Processed coverage track; coordinate conversion noted. Other processing not specified in supplied source table. |
| D. melanogaster | H3K4me3 | GSE20, GSE20787 | .bedGraph.gz | BDGP5 → BDGP6 | dm3ToDm6.over.chain.gz | Processed coverage track; coordinate conversion noted. Other processing not specified in supplied source table. |
| D. melanogaster | H3K9ac | GSE20, GSE20790 | .bedGraph.gz | BDGP5 → BDGP6 | dm3ToDm6.over.chain.gz | Processed coverage track; coordinate conversion noted. Other processing not specified in supplied source table. |
| D. melanogaster | H3K9me2 | GSE20, GSE20792 | .bedGraph.gz | BDGP5 → BDGP6 | dm3ToDm6.over.chain.gz | Processed coverage track; coordinate conversion noted. Other processing not specified in supplied source table. |
| D. melanogaster | H3K9me3 | GSE20, GSE20794 | .bedGraph.gz | BDGP5 → BDGP6 | dm3ToDm6.over.chain.gz | Processed coverage track; coordinate conversion noted. Other processing not specified in supplied source table. |
| D. melanogaster | H4K5ac | GSE20, GSE20800 | .bedGraph.gz | BDGP5 → BDGP6 | dm3ToDm6.over.chain.gz | Processed coverage track; coordinate conversion noted. Other processing not specified in supplied source table. |
| D. melanogaster | HP1 | GSE32, GSE32728 | .bedGraph.gz | BDGP5 → BDGP6 | dm3ToDm6.over.chain.gz | Processed coverage track; coordinate conversion noted. Other processing not specified in supplied source table. |
| D. melanogaster | Replication timing | GSM336, GSM336376 | .txt.gz | BDGP5 → BDGP6 | dm3ToDm6.over.chain.gz | Processed coverage track; coordinate conversion noted. Other processing not specified in supplied source table. |
| D. melanogaster | XR-seq (strand-specific) | GSM5600478/5661670 (S2); GSM5600476/5661668 (adult xpc KO); GSM5600452/5661644 (embryos) | .bw → .bedgraph | Not specified in supplied script/manifest; no liftOver performed | Not applied in supplied script | S2 (1 h CPD), adult xpc KO (2 h), embryo (2 h); 2 replicates per condition; bigWigs averaged per strand with bigwigCompare (25-bp bins), converted to bedGraph. No liftOver in supplied script. |
| **Species** | **Covariate** | **Source / accession** | **Original format** | **Genome build / conversion** | **LiftOver chain** | **Processing / replicates** |
| P. pacificus | ATAC | ERR2263975 / ERR2263983 | .bam | Elpaco → Elpaco | — | Two sorted BAM replicates; deepTools bamCoverage (CPM normalisation, 25-bp bins); bedtools unionbedg; mean of available replicate signals per interval; no liftOver. |
| P. pacificus | GC content | Calculated from reference sequence | Derived | Elpaco | — | No external dataset accession required; calculation method to be documented in Methods. |
| P. pacificus | Expression | Jung GRN-shiny; values_All_nottransformed.c-sv (GitHub) | .csv ->.tsv | PPA gene names mapped to Elpaco GTF gene IDs (WBGene); no coordinate liftOver | Not applicable | Embryonic time series; AG_<timepoint>_<replicate> columns grouped by timepoint; rowMeans across replicates (na.rm=TRUE); PPA-to-WBGene mapping from Elpaco genes.gtf; one Gene_ID/logVST TSV per timepoint. |
| P. pacificus | H3K27ac | ERR2263970 / ERR2263978 | .bam | Elpaco → Elpaco | — | Two sorted BAM replicates; deepTools bamCoverage (CPM normalisation, 25-bp bins); bedtools unionbedg; mean of available replicate signals per interval; no liftOver. |
| P. pacificus | H3K27me3 | ERR2263971 / ERR2263979 | .bam | Elpaco → Elpaco | — | Two sorted BAM replicates; deepTools bamCoverage (CPM normalisation, 25-bp bins); bedtools unionbedg; mean of available replicate signals per interval; no liftOver. |
| P. pacificus | H3K36me3 | ERR2263974 / ERR2263982 | .bam | Elpaco → Elpaco | — | Two sorted BAM replicates; deepTools bamCoverage (CPM normalisation, 25-bp bins); bedtools unionbedg; mean of available replicate signals per interval; no liftOver. |
| P. pacificus | H3K4me1 | ERR2263968 / ERR2263976 | .bam | Elpaco → Elpaco | — | Two sorted BAM replicates; deepTools bamCoverage (CPM normalisation, 25-bp bins); bedtools unionbedg; mean of available replicate signals per interval; no liftOver. |
| P. pacificus | H3K4me3 | ERR2263969 / ERR2263977 | .bam | Elpaco → Elpaco | — | Two sorted BAM replicates; deepTools bamCoverage (CPM normalisation, 25-bp bins); bedtools unionbedg; mean of available replicate signals per interval; no liftOver. |
| P. pacificus | H3K79me2 | ERR2263972 / ERR2263980 | .bam | Elpaco → Elpaco | — | Two sorted BAM replicates; deepTools bamCoverage (CPM normalisation, 25-bp bins); bedtools unionbedg; mean of available replicate signals per interval; no liftOver. |
| P. pacificus | H3K9me3 | ERR2263973 / ERR2263981 | .bam | Elpaco → Elpaco | — | Two sorted BAM replicates; deepTools bamCoverage (CPM normalisation, 25-bp bins); bedtools unionbedg; mean of available replicate signals per interval; no liftOver. |

